# Structure of the antidiuretic hormone vasopressin receptor signalling complex

**DOI:** 10.1101/2020.12.22.424028

**Authors:** Julien Bous, Hélène Orcel, Nicolas Floquet, Cédric Leyrat, Joséphine Lai-Kee-Him, Gérald Gaibelet, Aurélie Ancelin, Julie Saint-Paul, Stefano Trapani, Maxime Louet, Rémy Sounier, Hélène Déméné, Sébastien Granier, Patrick Bron, Bernard Mouillac

## Abstract

Arginine-vasopressin (AVP) is a neurohypophysial peptide known as the antidiuretic hormone. It forms an active signalling complex with the V2 receptor (V2R) and the Gs protein, promoting a cAMP/PKA-dependent aquaporin insertion in apical membranes of principal cells of the renal collecting ducts and ultimately, water reabsorption. Molecular mechanisms underlying activation of this critical G protein-coupled receptor (GPCR) signalling system are still unknown. To fill this gap of knowledge, we report here the structure of the AVP-V2R-Gs complex using cryo-electron microscopy (cryo-EM). Single-particle analysis revealed the presence of three different states. The two best maps were combined with computational and NMR spectroscopy constraints to reconstruct two structures of the ternary complex. These structures differ in AVP and Gs binding modes and could thus represent distinct complex conformations along the signalling activation pathway. Importantly, as compared to those of other class A GPCR-Gs complexes, the structures revealed an original receptor-Gs interface in which the Gsα subunit penetrates deeper into the active V2R, notably forming an ionic bond between its free C-terminal carboxylic function and the side chain of R137 in the V2R. Interestingly, the structures help to explain how V2R R137H or R137L/C variants can lead to two severe genetic diseases with opposite clinical outcomes, cNDI or NSIAD respectively. Our study thus provides important structural insights into the function of this clinically relevant GPCR signalling complex.

---

The biological actions of AVP, a cyclic nonapeptide, are mediated through three GPCR subtypes, Via, V1b and V2^1^. In addition, AVP is able to activate the related oxytocin (OT) receptor (OTR)^2^. The V2R is mainly expressed at the basolateral membrane of principal cells of the kidney collecting ducts and governs the crucial physiological function of body water homeostasis^3^. Binding of AVP to the V2R increases cAMP intracellular level via coupling to the adenylyl cyclase stimulatory Gs protein, leading to activation of protein kinase A, phosphorylation of aquaporin 2 water channels^4^ and, ultimately to water reabsorption and urine concentration. Activation of the V2R also elicits arrestindependent pathways such as receptor internalization and MAP kinase phosphorylation associated with cell growth and proliferation^5,6^. This GPCR is involved in many water balance disorders (hyponatremia consecutive to congestive heart failure, hypertension or hepatic cirrhosis) and voiding disorders (incontinence, nocturia), and as such constitutes a major therapeutic target^7^. Moreover, inactivating and constitutively active mutations in the V2R sequence are responsible for two rare X-linked genetic diseases with opposite clinical outcomes: 1) congenital nephrogenic diabetes insipidus (cNDI) characterized by excessive urine voiding^8^, 2) nephrogenic syndrome of inappropriate antidiuresis (NSIAD) characterized by excessive water loading and hyponatremia^9^. V2R is also a target for treating autosomal dominant polycystic kidney disease, the most frequent Mendelian inherited disorder affecting millions of people worldwide^10^. This pathology results from increased cell proliferation, apoptosis and dedifferentiation, in which cAMP- and MAP kinase-dependent signaling pathways are highly activated.

The structural biology of GPCRs has made significant progress during the last decade with a wealth of information on ligand binding and G protein coupling that shed light on structural and dynamic aspects of their function^11,12^. Yet V2R like many GPCRs, has been refractory to high-resolution structure determination. Cryo-EM has emerged as a powerful method for the determination of challenging membrane protein structures^13^, in particular when the intrinsic structural dynamics of the target prevents the use of crystallogenesis. A growing list of GPCR-G protein complex structures has thus been determined^14,15^, revealing key molecular mechanisms of agonist binding and G protein (Gi, Gs, Gq, Go) coupling to class A and class B GPCRs. Here we have developed an *in vitro* purification strategy to reconstitute the GPCR signalling complex comprising the AVP-bound V2R and the heterotrimeric Gs protein stabilized with the nanobody Nb35. Cryo-EM single-particle analysis revealed the presence of three distinct populations of the ternary complex with two best maps at a mean resolution of 4.0 and 4.1 Å. A novel hybrid approach was used to build both corresponding structures. Analyses of the structural features of the distinct conformations provide unprecedented molecular insights into the dynamic process of ternary complex formation between the hormone AVP, the V2 receptor and the Gs protein.

## Determination of the AVP-V2R-Gs-Nb35 complex structure

After extensive biochemical optimization, validation of the complex preparation and of cryo-EM grid sample vitrification, (see Methods, **Extended Data Fig. 1**, **Supplementary Figs. 1-2**), a total number of 25,770 movies were recorded, with 3.5 million particles picked and sorted out for further data processing (**Extended data Fig. 2-3**). After 3D classification of projections and 3D refinement, we identified three different conformational states of the complex, referred to as Loose (L), Tight-1 (T1) and Tight-2 (T2). Reconstruction of each state was at 4.2 Å, 4.5 Å and 4.7 Å, with a distribution of 16%, 48% and 36%, respectively (**Extended Data Fig. 2**), the local resolution varying from 3.2 to 6.4 Å (**Extended data Fig. 3c-d).** Using the recent algorithm developed to enhance cryo-EM maps by density modification^16^, the resolution of density maps were improved to 4.0 (L state), 4.1 (T1 state) and 4.5 Å (T2 state), respectively (**Extended data Fig. 2, Extended Data Table 1, Supplementary Fig. 3)**. These maps mainly differ in the angle of Gs-Nb35 with the receptor 7TM, and may reflect an inherent high flexibility of the complex. A conformational heterogeneity analysis using multi-body refinement revealed that more than 78% of the variance is accounted for by the fourth first eigenvectors related to rotations and translations between AVP-V2R and Gs-Nb35 (**Extended Data Fig. 4**, **Extended data Movie 1**). The 4.5 Å map of the T2 state was not well enough resolved to compute a reliable structure. Therefore, only the L and T1 structures, referred to as L and T states, were used for further analysis.

Because we could not unambiguously build the AVP in the calculated maps, we designed an original hybrid strategy based on a combination of cryo-EM maps, computational molecular dynamic simulations and experimental saturation transfer difference (STD) NMR (**Fig. 1a-e**, **Supplementary Discussion and Figs. 4-8**). Based on this approach, the L and T models were then built in a more conventional manner to match as closely as possible the density maps (**Fig. 1f-g, Extended Data Table 1**).

**Fig. 1.**
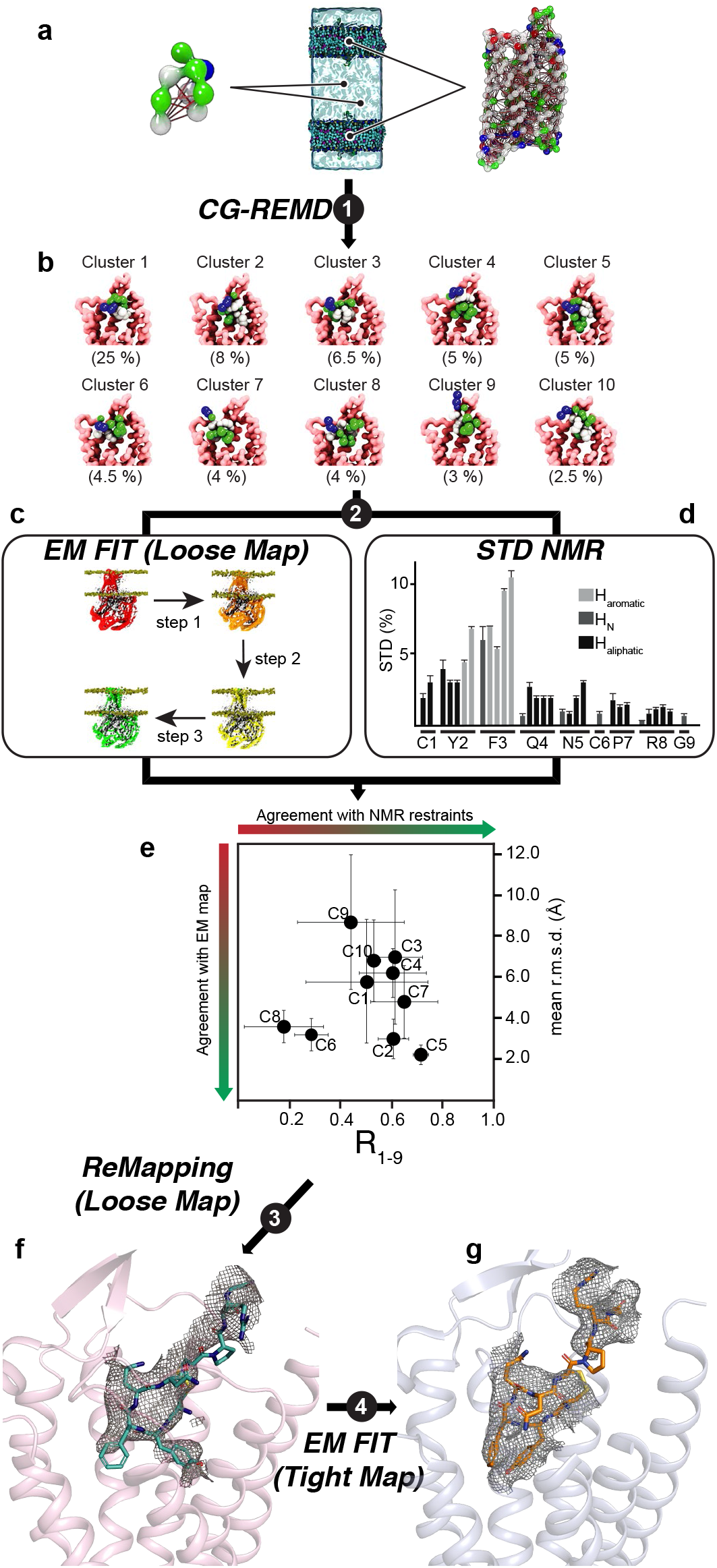
Overview of the hybrid strategy: a combination of cryo-EM, computational and NMR. **a**) schematic representation of the unbiased CG ab-initio approach. The internal elastic networks used for both the AVP (left) and the V2R (right) are shown. The full system (centre) used for the CG-REMD simulations included two receptors and two peptides in order to create an artificial extracellular compartment and improve the conformational sampling. **b**) 10 most populated clusters obtained for the AVP-V2R complex after the 3 independent CG-REMD simulations. The upper parts of TM 6 and TM7 of V2R were removed for clarity. The 10 clusters describe 67.5% of the whole conformations analysed in the procedure. **c**) schematic representation of the successive steps employing the CDMD method to fit the models resulting from the CG-REMD simulations into the L cryo-EM map of the AVP-V2R-Gs-Nb35 complex. **d**) Mapping AVP contact surface by experimental STD NMR. The STD effect profile (in %) is shown as a function of AVP protons (aliphatic, N (backbone), aromatic). Intense STD signals were only observed in the presence of V2R, mostly for the aromatic protons of Tyr^2^ and Phe^3^ (and to a lesser extent of Cys^1^) residues of AVP. **e**) Cross-correlation between computational and STD NMR. First, variability of the position of AVP was calculated as mean RMSD values (in Å) after cross-comparison of models (5 for each cluster) resulting from the fitting procedure of each of the 10 clusters in the L density map. Cluster 5 showed the smaller variability (2.2 Å). Second, experimental STD values were compared to the expected STD values from all-atom models issued from MD simulations and correlation coefficients were calculated for the whole peptide (R_1-9_). Cluster 5 appeared as the best cluster fitting to experimental STD values. **f**) Remapping into the L density map. Based on cluster 5, the final L model was then built in a more conventional manner to match as closely as possible the L density map. **g**) Based on the L model, the T model was built to match the T density map.

In the final models, side chains of most residues are clearly identifiable in the 7 TM and helix 8 of the V2R in both structures (**Extended Data Fig. 5**). Intracellular loop1 (ICL1) was well defined in the maps, as well as the contacts between V2R and the Gs protein. The α-helical domain of Gαs subunit was subtracted during single particle analysis for high-resolution map refinement. Intracellular loops 2 and 3 and the C-terminus of V2R were not seen in the density maps, and were not constructed in the final models.

## Overall architecture of the ternary complex

Both L and T AVP-V2-Gs ternary complexes present a typical GPCR-G protein architecture with the receptor 7TM helix bundle engaging the peptide agonist on the extracellular side and the Gαs C-terminus domain (α5 helix) on the intracellular side (**Fig 2a-d**). However, the L and T states present large structural differences most notably in the position of the G protein heterotrimer relative to V2R (**Fig. 2e**). The α5 helix interacts more tightly in the T state than in the L state (**Fig. 2b and 2d**), inducing a translation of the whole Gs heterotrimer (**Fig. 2e**). In particular, the α4 helix and the Ras-like domain of Gαs are translated from 4 Å and 5 Å between the L and T states, respectively. These movements position the αN helix 5 Å closer to the receptor in the T state in comparison to the L state (**Fig. 2e**). Those Gα movements are also accompanied by a 7 Å translation of the Gβ N-terminal helix, a translation of the γ subunit of 6 Å and a translation of Nb35 of 7 Å (**Fig. 2e**).

**Fig. 2.**
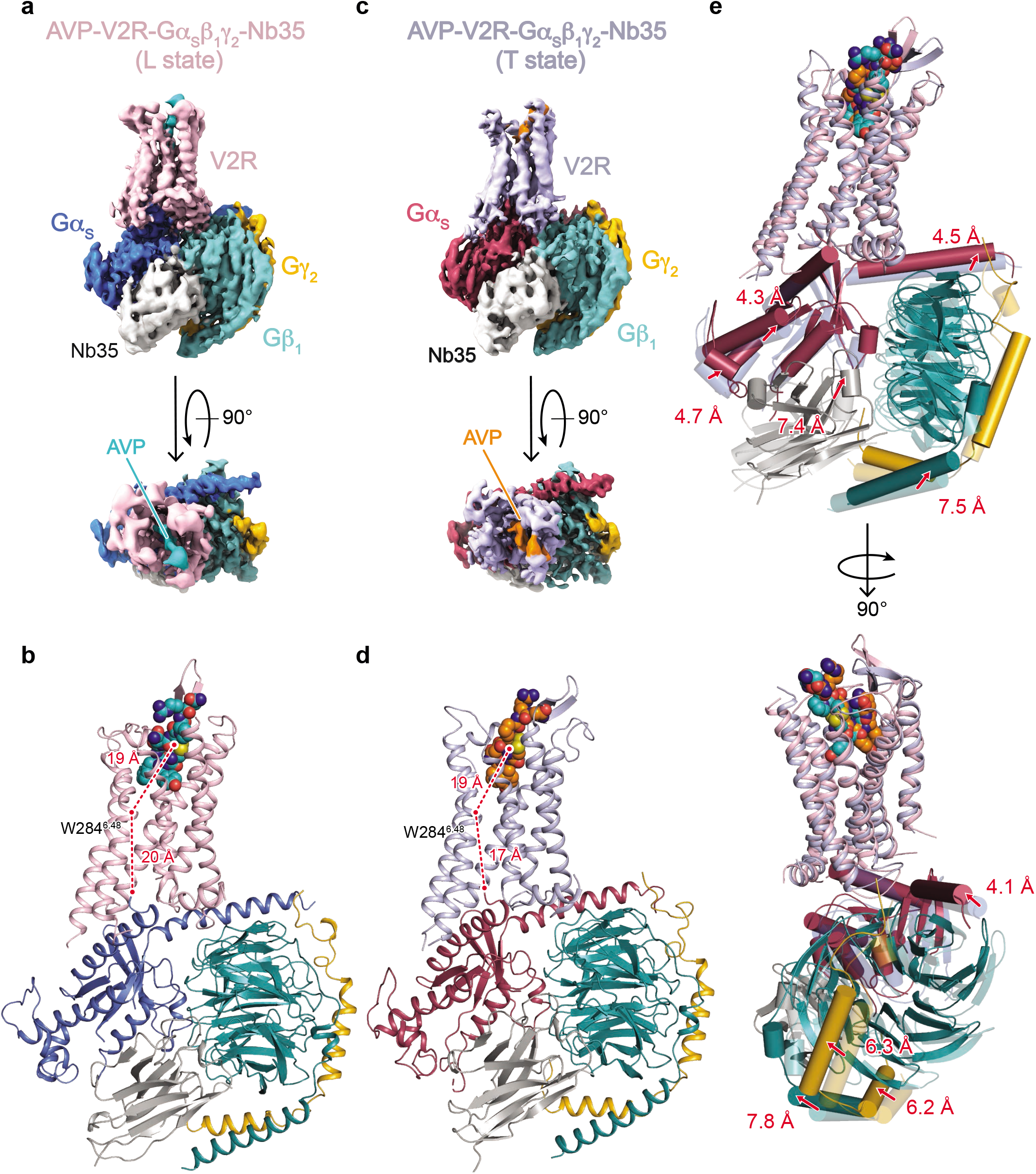
Structures of AVP-V2R-Gs-Nb35 complexes in L and T conformations. **a**) orthogonal views of the cryoEM density maps of the L state of the AVP-V2R-Gs-Nb35 complex and **b**) corresponding model as ribbon representation. V2R is coloured in pink, Gαs in dark blue, G 1 in turquoise, Gγ2 in yellow, Nb35 in grey, AVP in cyan. In **b**, the distances between W284^6.48^ (at its Cα carbon) and the AVP center of mass, and between W284^6.48^ and the C-terminal end of 5 helix of Gs (at the Cα carbon of the free carboxylic acid) are shown. **c**) and **d**) corresponding maps and model for the T state. V2R is coloured in blue grey, Gαs in raspberry, Gβ1 in turquoise, Gγ2 in yellow, Nb35 in grey, AVP in orange. In **d**, distances are measured as in **b**. **e**) L and T models are aligned on the V2R chains, and rotations/translations are shown by measuring displacement (in Å) of Gαs, Gβ1, Gγ2 and Nb35.

The presence of several conformational states and the multi-body refinement analysis reflect the dynamics of V2-Gs complex formation. From the final L structure model, a principal component analysis (PCA) obtained from classical molecular dynamic simulations revealed similar dynamics and suggests that the conformations captured by the cryo-EM 3D reconstructions represent averaged states that are part of a much larger conformational ensemble (**Extended data Movie 2, Supplementary Fig. 9**). Though those differences are less pronounced that the ones recently described for the neurotensin receptor NTSR1-Gi1 complexes^17^, they further indicate that GPCR-G protein coupling is a dynamic process in which the G protein may explore different set of conformations.

## AVP binding pocket within V2R and comparison with OTR binding site

Our hybrid approach allowed us to build convincing models of AVP binding poses in both L and T structures. The final calculated structures present a central position of AVP in the orthosteric pocket of the V2R along the axis of the helical bundle **(Fig. 3, Extended Data Fig. 6)**. The extracellular domains of the V2R are widely opened in both L and T conformations, a feature consistent with the accommodation of a cyclic peptide like AVP (**Extended Data Fig. 6**), and in agreement with the recently reported inactive OTR structure^18^. In the L and T structures, AVP contacts residues from both TM helices and extracellular loops **(Fig. 3a-c, Extended Data Fig. 6**) in agreement with what was originally proposed based on pharmacological data^19^. Consistent with its amphipathic nature, AVP interacts with two chemically distinct interfaces in a 15 Å deep binding pocket to form both polar and hydrophobic contacts **(Fig. 3a-c)**.

**Fig. 3.**
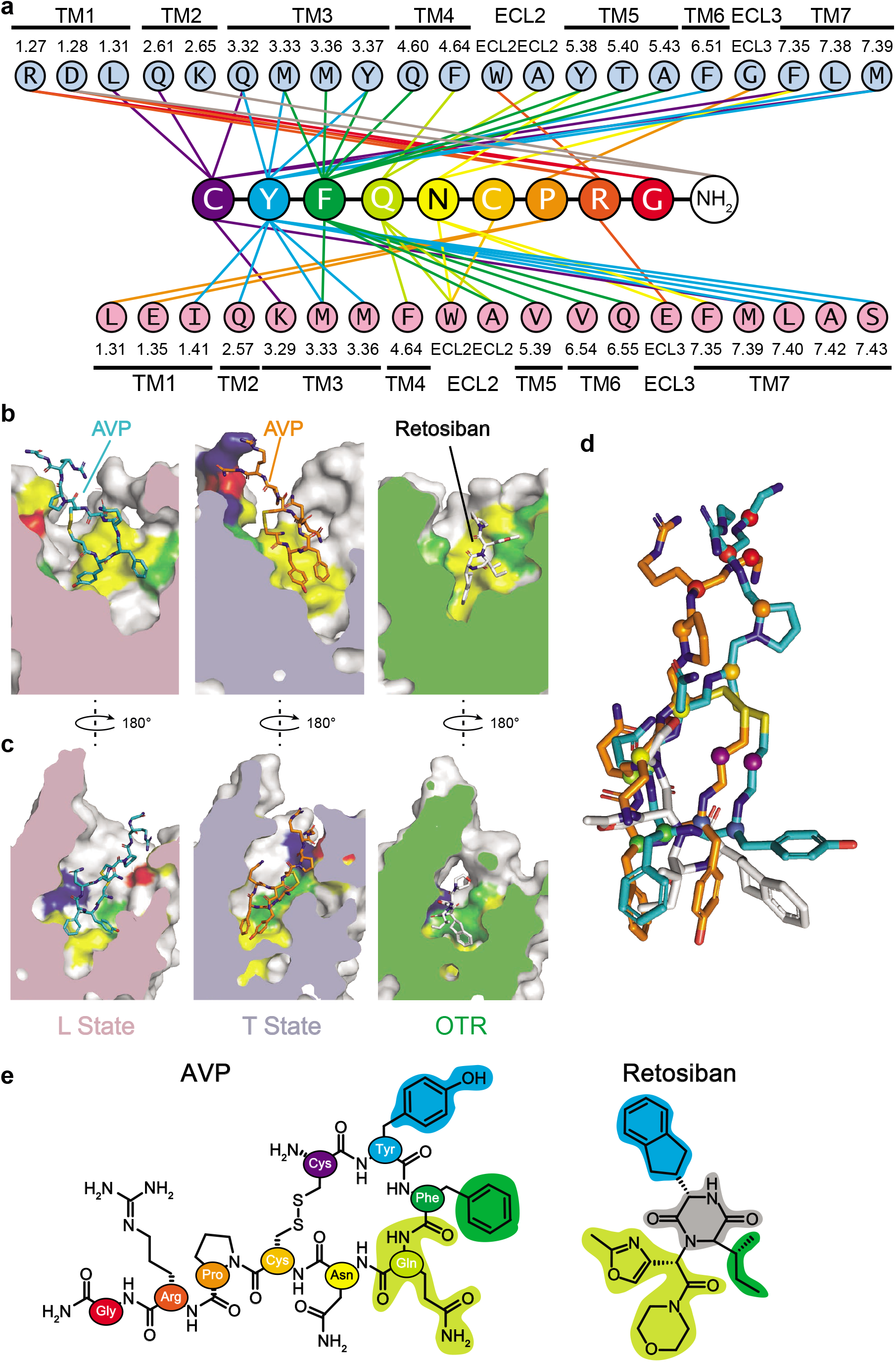
AVP binding site of the V2R, comparison with retosiban binding site in OTR. **a**) Direct contacts between AVP and V2R in L and T structures. Interactions (within a maximum 5 Ä distance) are shown between each AVP residue (and the C-terminal amide) with V2R residues in the L structure (pink) and in the T structure (blue). All TM helices and extracellular loop 2 and 3 interact with the hormone AVP. V2R residues are labeled according to the Ballesteros-Weinstein numbering. Each residue from AVP is colored differently for clarity. **b**) Side views of the binding pocket in the L and T structures and in the inactive structure of OTR. AVP binding modes in the L (pink) and T (light blue) structures, are compared to that of the small molecule antagonist retosiban in the OTR structure (green), all viewed from TM3. Residues from receptors that interact with AVP or retosiban are depicted in different colors: yellow for hydrophobic, green for polar, red and dark blue for negatively and positively charged, respectively. **c**) Side views of the binding pockets after a 180° rotation. AVP and retosiban are viewed from TM6. The same color code is used. **d**) Superimposition of AVP and retosiban. The peptide agonist and the nonpeptide antagonist are superimposed after alignment of V2R and OTR structures. The most hydrophobic parts of both ligands superimpose at the bottom of the orthosteric binding pocket. **e**) Structure comparison of AVP and retosiban. AVP is shown using the same color code than in pannel a. The retosiban indanyl moiety, the sec-butyl group, and the oxazol-morpholine amide moiety superimpose with AVP Tyr2, Phe3 and Gln4 respectively. The retosiban 2,5-diketopiperazine core is positioned between AVP Tyr2 and Phe3.

While AVP conformations occupy a central position in both the L and T binding clefts, interesting changes are observed due to a translation of the Tyr^2^ residue side-chain (from TM7 to TM3), and to a movement of the C-terminal tripeptide (inversion in Arg^8^ and Gly^9^-NH2 positions) at the V2R surface (**Extended data Fig. 6**). The cyclic part of AVP (Cys^1^ to Cys^6^) and the Pro^7^ are buried into the cleft defined by the seven-helix bundle of V2R, leaving only Arg^8^ residue and C-terminal glycinamide exposed to the solvent (**Extended data Fig. 6).** In both the L and T structures, the Cys^1^-Tyr^2^-Phe^3^ hydrophobic motif of AVP binds deeper in the binding site creating key contacts with the receptor (**Fig. 3a-c**), in agreement with STD spectroscopy data (**Fig. 1**).

V2R and OTR belong to the same subfamily of peptide class A GPCRs and share a common orthosteric binding site^19, 20^. Although V2R and OTR (PDB 6TPK) structures^18^ represent different GPCR conformations (active agonist-bound V2R *vs* inactive antagonist-bound OTR), it is interesting to compare the complete set of residues involved in the binding of the natural hormone AVP with the ones involved in retosiban binding to gain insights into ligand binding and efficacy in this receptor family **(Fig 3b-e).** Many OTR residues involved in the binding of retosiban are actually conserved among AVP/OT receptors and also interact with AVP in the V2R **(Fig. 3b-e).** Interestingly, the conserved W^6.48^ and F^6.51^ (Ballesteros-Weinstein numbering) in AVP/OT receptors, interact with the highly hydrophobic indanyl moiety of retosiban in the crystal structure of inactive OTR. AVP also makes contact with F^6.51^ through its Tyr^2^ but is not in contact with W^6.48^ in the V2R, probably because it is too bulky to bind deeper in the pocket. These data confirm that hydrophobic small molecule nonpeptide antagonists and AVP partially superimpose at the bottom of the orthosteric binding pocket of AVP/OT receptors **(Fig. 3d-e)**^21,22,23^.

## Activation of the V2R and comparison with other class A GPCRs

The active-state structures of the V2R reveal key structural features of the activation process by comparison with the OTR inactive structure (**Fig. 4a-e**). Moreover, to get a more general view of V2R activation, it was also important to look at the canonical conformational changes of TMs and of conserved motifs involved in other ligand-activated GPCRs of class A^24,25^. Thus, compared to other active GPCR structures and to the inactive antagonist-bound OTR structure (**Fig. 4, Supplementary Fig. 10**), the L and T structures of V2R present all the features of active conformations, i.e. a large-scale displacement of TM6 (**Fig. 4a)**, conformational changes of W^6.48^ toggle switch **(Fig. 4b**), a rearrangement of the P^5.50^-S^3.40^-Y^6.44^ transmission switch, equivalent to the PIF motif in other GPCRs (**Fig. 4c**), a rotation of the conserved NPxxY^7.53^ motif **(Fig. 4d)** and a broken D136^3.49^-R137^3.50^ ionic lock (**Fig 4e, Fig. 5e**).

**Fig. 4.**
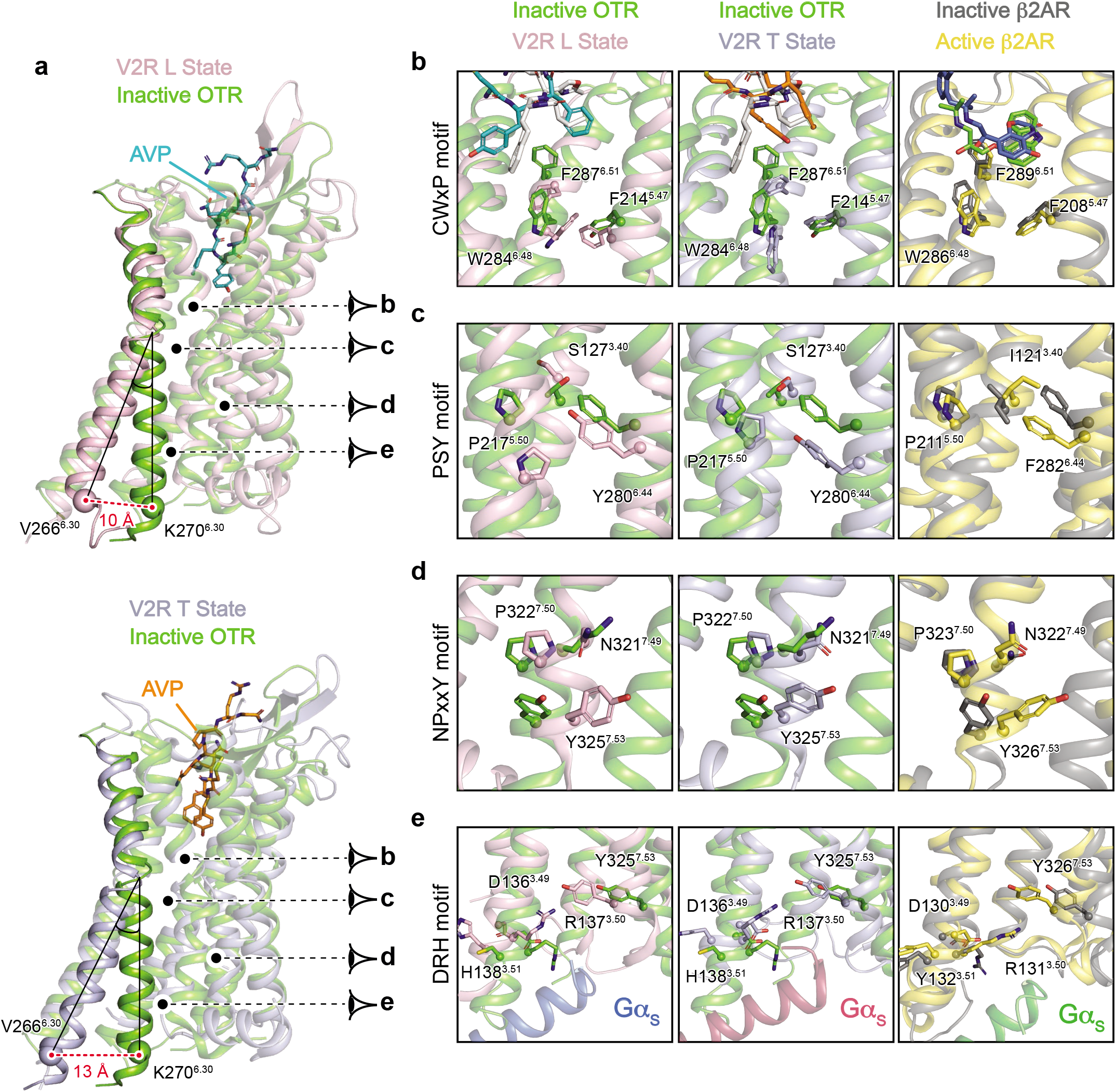
Active conformations of L and T V2R states, comparison with inactive structure of OTR and active/inactive structures of β2AR. **a**) Large-scale displacement of TM6. The V2R L (pink) and T (blue grey) active structures are aligned onto that of the inactive OTR (green) structure. Residue 6.30 (Ballesteros-Weinstein numbering) is chosen as a reference (V266 in V2R, K270 in OTR) for measuring the outward movement of TM6: 10 and 13 Å between OTR and V2R in the L and T states, respectively. Activation of molecular switches along the helix bundle of the V2R are viewed in panels b, c, d and e. For comparison, rearrangements of those corresponding motifs in the β2AR are depicted. **b**) Rotameric toggle switch in the conserved CWxP motif. Position of W^648^, F^6.51^ and F^5.47^ (284, 287 and 214 in V2R) are shown. **c**) Rearrangement of the PSY transmission switch. The P^5.50^-S^3.40^-Y^6.44^ motif (217, 127 and 280 in V2R) is equivalent to the PIF motif in other GPCRs. **d**) Rotation of the NPxxY conserved motif in TM7. The conserved Y^7.53^ (position 325 in V2R) is shown. **e**) Breaking of the conserved ionic lock in TM3. Upon activation of V2R, the ionic bond between D136^3.49^ and R137^3.50^ is broken and R137^3.50^ projects to Y325^7.53^.

**Fig. 5.**
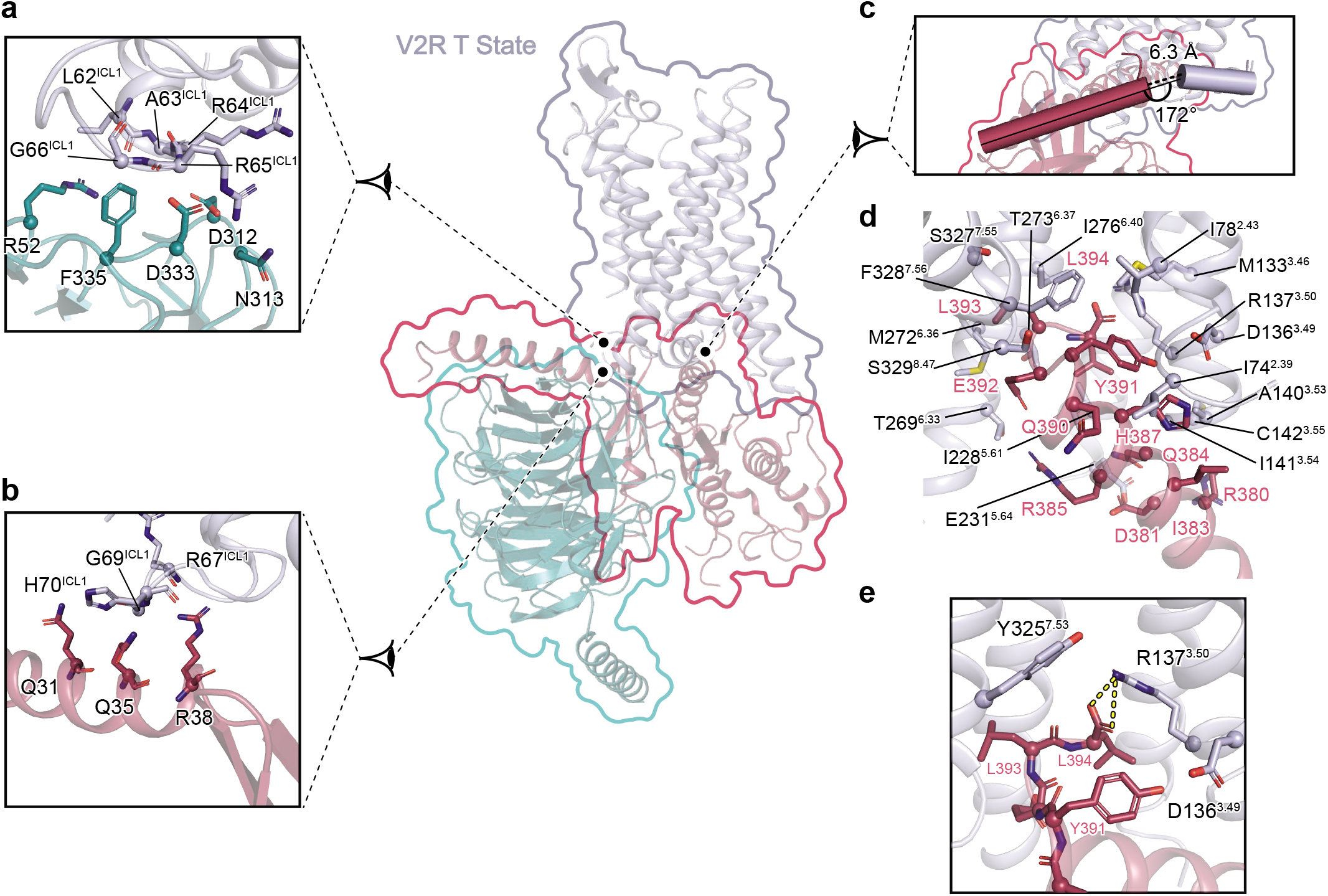
Interface of the V2R T state with Gs. Interactions between V2R and Gs heterotrimer are shown. Specific interfaces are depicted, and residues in close proximity (within a maximal 3.9 Å distance) are highlighted (blue grey for V2R, turquoise for Gβ subunit, raspberry for *Gα* subunit). **a**) Interaction of V2R ICL1 with G subunit. **b**) Interaction of V2R ICL1 with Nterminal helix of Gα subunit. **c**) Position of C-terminal h5 helix of Gα subunit relative to V2R helix 8. The distance between Gα and helix 8 is indicated. Angle between these two domains is shown. **d**) Interacting residues between the C-terminal h5 helix of Gα subunit and V2R. **e**) Zoom on an ionic bridge between the C-terminal free carboxylic moiety of h5 helix of Gα subunit and V2R R137^3.50^.

Importantly by comparing the structures of the inactive antagonist-bound OTR with the active agonist-bound V2R, it appears that contacts between M123^3.36^ and F287^6.51^-W284^6.48^ motif (all in contact with Tyr^2^ of AVP) undergo large conformational rearrangements (**Extended Data Fig. 6**). It is thus tempting to speculate that it is a key motif regulating the activity of this family of receptor.

As indicated above, the V2R R137^3.50^ participates in the ionic lock motif involved in the balance of active *versus* inactive states of class A GPCRs^24^. In the inactive structure of OTR (**Fig. 4e**), D136^3.49^ and R137^3.50^ interact with each other through this ionic lock. For comparison, this salt bridge is broken in the L and T active conformations of the V2R-Gs complex (**Fig. 4e, Fig. 5e**). The observed constitutive activity towards Gs coupling for the missense mutations C137^3.50^ or L137^3.50^ responsible for NSIAD^9,26,27^ can thus be explained from a structural point of view since these hydrophobic residues are not able to form such an ionic lock to stabilize the inactive state. On the contrary, the mutant H137^3.50^ causing cNDI^28, 29^ might still be able to maintain the balance between active and inactive states of the V2R through its partial positive charge. Its loss-of-function rather reflects the loss of accessibility to AVP because of a constitutive internalization^27, 28, 29^.

## V2R-Gs interactions

The cryo-EM maps of the ternary complex clearly establish the structural details of V2R-Gs coupling. As anticipated from the conserved mechanism of GPCR-G protein coupling^30,31^, both the L and T conformations show a similar overall architecture of the complex interface with the engagement of the Gαs C-terminal α5 helix in the core of the 7TM **(Fig. 5, Extended Data Fig. 7, Supplementary Fig. 10)**. However, there are some interesting differences compared to other GPCR-Gs complex structures. Of note, in both the L and T structures, the V2R ILC1 makes many direct contacts with the Gβ subunit. In the T state, ICL1 residues L62-A63-R64-R65-G66 interact with Gβ R52, D312-N313 and D333-F335 (**Fig5a**). In the L state, ICL1 residues R65-G66-R67-R68 interact with Gβ R52, D312 and D333 (**Extended Data Fig. 7a**). These contacts between V2R and Gβ are much more numerous than in the class A GPCR β2AR- or A2AR-GS complexes^32, 33^. Moreover in the T conformation, there are some additional contacts between V2R ICL1 (R67-G69-H70) with the N-terminal α helix of Gαs (Q31, Q35 and R38), resulting in a more compact interaction (**Fig 5b**). In the L state, V2R (W71) and N-terminal α helix of Gαs (Q35 and R38) contacts are more limited (**Extended Data Fig. 7b**). Contacts between the N-terminal α helix of Gαs with GPCRs have only been seen in GLP1R and CTR class B GPCR complexes^34, 35^, not in class A GPCR-G complexes.

In contrast to what was observed for the β2AR^32^ and the μOR^36^, the Gαs C-terminal α5 helix appears to extend helix 8 (H8) of the V2 receptor, lying almost parallel to the membrane plane **(Fig. 5c, Extended Data Fig 7c, f-h)**. Also, compared with the β2AR, the C-terminus of Gαs is interacting deeper in the V2R 7TM core making direct contact with residues (L and T states, respectively) of V2R that are part or in close proximity to the conserved NPxxY (TM7) and DRH (TM3) activation motifs **(Fig 5d-e, Extended Data Fig. 7d-h)**. In this respect, the V2-Gs interaction resembles more the interaction seen in the μOR-Gi complex **(Extended Data Fig. 7f-h)**. The V2R TM7-H8 hinge region also makes a strong contact with the Gαs ELL motif, in particular through hydrophobic contacts with the F328^7.56^ side chain **(Fig. 5d)**. The T and L conformations differ here in the position of the Gαs L394 side chain originating from a distinct F328^7.56^ side chain conformation (pointing towards I78^2.43^ of the receptor in the T structure or towards Gαs L394 in the L structure) (**Fig. 5d, Extended Data Fig. 7d**). Most notably in the T state, the side chain of R137^3.50^ which is part of the ionic lock motif, forms an ionic interaction with the free carboxylic acid function of the Gαs C-terminus (**Fig. 5e**), a direct contact that was not observed before between a GPCR and a G protein of any family (Gs, Gi, Go or Gq)^14,15,37^. Moreover, in the L state, the density map suggests that the R137^3.50^ side chain could adopt two conformations, one forming a similar ionic interaction with the carboxylic acid of Gαs L394 main chain, and, the other one, pointing towards the Y325^7.53^ from the NPxxY motif (**Extended Data Fig. 7e).**

## Conclusion

In this study, we solved two structures of the active state of the AVP hormone-bound V2R in complex with the Gs protein. The structural insights into the function of this signalling system pave the way for future drug development to treat water balance disorders^7^.

## METHODS

### Data analysis and Figure preparation

Figures were created using the PyMOL 2.3.5 Molecular Graphics System (Schrödinger, LLC) and the UCSF Chimera X 0.9 package. Data were plotted with GraphPad Prism 8.3.0.

### V2R expression and purification

The version of the human V2R used for this study in described in Supplementary Methods and was cloned into a pFastBac™1 vector. It was expressed in Sf9 insect cells using the Bac-to-Bac® baculovirus expression system ((Thermofisher) according to manufacturer’s instructions. Insect cells were grown in suspension in EX-CELL® 420 medium (SigmaAldrich) to a density of 4 × 10^6^ cells per ml and infected with the recombinant baculovirus at a MOI of 2-3. The culture medium was supplemented with the V2R pharmacochaperone antagonist Tolvaptan (TVP, SigmaAldrich) at 1 μM to increase the receptor expression levels^38, 39^. The cells were infected for 48h-54h at 28°C and expression of the V2R was checked by immunofluorescence using an anti-Flag M1 antibody coupled to Alexa488. Cells were then harvested by centrifugation (2 steps, 20 min, 3,000xg) and pellets were stored at −80°C until use.

The cell pellets were thawed and lysed by osmotic shock in 10 mM Tris-HCl pH 8, 1 mM EDTA buffer containing 2mg.ml^−1^ iodoacetamide, Tolvaptan 1 μM and protease inhibitors (leupeptine 5μg.ml^−1^, benzamidine 10μg.ml^−1^ and PMSF 10μg.ml^−1^). After centrifugation (15 min, 38,400xg), the pellet containing crude membranes was solubilized using a glass dounce tissue grinder (15 + 20 strokes using A and B pestles, respectively) in a solubilization buffer containing 20 mM Tris-HCl pH 8, 500 mM NaCl, 0.5 % (w/v) n-dodecyl-β-D-maltopyranoside (DDM, Anatrace), 0.2% (w/v) sodium cholate (SigmaAldrich), 0.03 % (w/v) cholesteryl hemisuccinate (CHS, SigmaAldrich), 20 *%* Glycerol, 2 mg.ml^−1^ iodoacetamide, 0.75 ml.L^−1^ Biotin Biolock (IBA), Tolvaptan 1 μM and protease inhibitors. The extraction mixture was stirred for 1 h at 4 °C and centrifuged (20 min, 38,400xg). The cleared supernatant was poured onto an equilibrated Strep-Tactin® resin (IBA) for a first affinity purification step. After 2h of incubation at 4°C under stirring, the resin was washed three times with 10 CV of a buffer containing 20 mM Tris-HCl pH 8, 500 mM NaCl, 0.1 % (w/v) DDM, 0.02% (w/v) sodium cholate, 0.03 % (w/v) CHS, Tolvaptan 1 μM. The bound receptor was eluted in the same buffer supplemented with 2.5 mM desthiobiotin (IBA).

The eluate was supplemented with 2mM CaCl_2_ and loaded onto a M1 anti-Flag affinity resin (SigmaAldrich). The resin was washed with 10 CV of two successive buffers containing 20 mM HEPES pH 7.5, 100 mM NaCl, 0,1% DDM, 0,01% CHS, 10 μM AVP, 2mM CaCl_2_, then 20 mM HEPES pH 7.5, 100 mM NaCl, 0,025% DDM, 0,005% CHS, 10 μM AVP, 2mM CaCl_2_, respectively. The receptor was eluted from the Flag resin using a buffer containing 20 mM HEPES pH 7.5, 100 mM NaCl, 0,025% DDM, 0,005% CHS, 10 μM AVP, 2 mM EDTA and 200 μg/mL Flag peptide (Covalab).

After concentration using a 50kDa MWCO concentrator (Millipore), the V2R was purified by size exclusion chromatography (SEC) using a Superdex 200 (10/300 column) connected to an AKTA purifier system (GE Healthcare). Fractions corresponding to the pure monomeric receptor were pooled (~ 2 ml) and concentrated to 50-100 μM with an excess of AVP (200 μM).

### Gs expression and purification

Human Gαs, Gβ1 with a N-terminal Twin-Strep-tag® and Gγ2 were all expressed in Sf9 insect cells grown in EX-CELL® 420 medium (SigmaAldrich). A recombinant baculovirus for Gαs subunit was prepared using the BestBac™ (Expression Systems) strategy, whereas a baculovirus for Gβ1 and Gγ2 was prepared using the Bac-to-Bac® system. Gβ1 and Gγ2 were cloned in tandem into the pFastBac™ Dual vector (Thermofisher). Sf9 cells, at a density of 4×10^6^ cells/ml, were co-infected with both viruses at a 1:2 Gαs:Gβ1γ2 ratio for 72h at 28°C. Cells were harvested and pellets stored at −80°C.

Co-infected Sf9 cell pellets were thawed and lysed in a buffer containing 10 mM TRIS pH 7.4, 1mM EDTA, 5 mM β-Mercapto-Ethanol, 10 μM GDP and protease inhibitors (leupeptine 5μg.ml^−1^, benzamidine 10μg.ml^−1^ and PMSF 10μg.ml^−1^). Lysed cells were centrifuged (20 minutes, 38,400xg). The pellets containing the crude membranes were homogeneized using a glass dounce tissue grinder (20 strokes with tight B pestle) in solubilization buffer containing 20 mM HEPES pH 7.5, 100 mM NaCl, 1% DDM, 5 mM MgCl_2_ supplemented with 5 mM β-Mercapto-Ethanol, 10 μM GDP, 0.75 ml.L^−1^ biotin biolock and protease inhibitors. The mixture was stirred for 40 minutes at 4°C and centrifuged (20 min, 38,400xg). The supernatant was loaded onto a Strep-Tactin® affinity resin equilibrated with the same buffer. The resin was washed 3 times, first with 5 CV of solubilization buffer, then with 5 CV of solubilisation buffer supplemented with 100 μM TCEP (instead of β-Mercapto-Ethanol), and finally with 10 CV of wash buffer containing 20 mM HEPES pH 7.5, 50 mM NaCl, 0.1% DDM, 1 mM MgCl_2_, 100 μM TCEP, 10 μM GDP. The Gs heterotrimer protein was eluted in the same buffer supplemented with 2.5 mM desthiobiotin. After a treatment with antarctic phosphatase (5 Units, NEB Inc.) for 30 minutes at 4°C, the Gs protein was concentrated to 10 mg/ml using 50 kDa MWCO concentrators. 20% glycerol was added to the sample and aliquots were flash-frozen in liquid nitrogen before storage at −80 °C.

### Nb35 expression and purification

The production and purification of Nb35 were performed following a protocol established by Kobilka and coworkers^40^. It is described in Supplementary Methods.

### Purification of the AVP-V2R-Gs-Nb35 complex

Formation of a stable complex was done by mixing the purified V2R with 1.2 molar excess of purified Gs heterotrimer, 250 μM AVP and 2.5 mM MgCl_2_. The coupling reaction was allowed to proceed at RT for 45 minutes and was followed by addition of apyrase (0.0125 Units, NEB Inc.) to hydrolyse residual GDP and maintain the high-affinity nucleotide-free state of Gs. 15 minutes later, Nb35 was added at a 2-fold molar excess compared to Gs. After 15 more minutes at RT, the mix was incubated overnight at 4°C. In most reaction mixtures, the final concentration of V2R was 20-30 μM, that of Gs 30-40 μM and the one of Nb35 around 80 μM. To remove excess of G protein heterotrimer and Nb35, the complex AVP-V2R-Gs-Nb35 was purified by a M1 anti-Flag affinity chromatography. After loading, the DDM detergent was then gradually exchanged with Lauryl Maltose Neopentyl Glycol (LMNG, Anatrace). The LMNG concentration was then decreased gradually from 0.5% to 0.01%. The complex and the unbound V2R were eluted in 20 mM HEPES pH 7.5, 100 mM NaCl, 0.01% LMNG, 0.002% CHS, 2 mM EDTA, 10 μM AVP and 0.2 mg.mL^−1^ Flag peptide. The eluted AVP-V2R-Gs-Nb35 complex was separated from unbound V2R by SEC on a Superdex 200 10/300 with a buffer containing 20 mM HEPES pH 7.5, 100 mM NaCl, 0.002% LMNG, 0.0025% glyco-diosgenin (GDN, Anatrace), 0.002% CHS, 10 μM AVP. The fractions corresponding to the complex were collected, concentrated with a 50 kDa MWCO concentrator and subjected to a second SEC on a Superose 6 (10/300 GL, GE Healthcare) with a buffer containing 20 mM HEPES pH 7.5, 100 mM NaCl, 0.0011% LMNG, 0.001% GDN, 0.002% CHS, 10 μM AVP. Peak fractions were pooled and concentrated using a 50 kDa MWCO concentrator to concentrations ranging from ~1 mg.mL^−1^ to ~4 mg.mL^−1^ for Cryo-EM studies. The amphipol A8-35 (Anatrace) was added at 0.001% to help in the dispersion of the particles for Cryo-EM grid preparation.

### Negative stain microscopy observations

Before preparing Cryo-EM grids, we first checked the quality and the homogeneity of the AVP-V2R-Gs-Nb35 sample by NS-EM. The procedure is described in Supplementary Methods.

### Cryo-EM sample preparation, image acquisition

In this study, two datasets have been recorded from two different preparations of AVP-V2R-Gs-Nb35. For the first dataset acquisition, 3 μL of purified AVP-V2R-Gs-Nb35 at 0.75 mg/ml concentration were applied on glow-discharged Quantifoil R 1.2/1.3 300 mesh copper holey-carbon grids (Quantifoil Micro tools GmbH, Germany), blotted for 4.5s and then flash-frozen in liquid ethane using the semi-automated plunge freezing device Vitrobot Mark IV (ThermoFisher) maintained at 100% relative humidity and 4°C. For the second dataset acquisition, cryo-EM grids were prepared as previously, but the purified V2R-Gs-Nb35 complex was at a concentration of 4 mg/ml and the cryo-EM grids were prepared using a EM GP2 (Leica microsystems) plunge freezer with a 4s blotting time (100% humidity and 4°C).

Images were collected in 2 independent sessions on a TEI Titan Krios (ThermoFisher) at the EMBL of Heidelberg (Germany) at 300 keV through a Gatan Quantum 967 LS energy filter using a 20 eV slit width in zero-loss mode and equipped with a K2-Summit (Gatan inc.) direct electron detector configured in counting mode. Movies were recorded at a nominal EFTEM (energy-filtered transmission electron microscope) magnification of ×165,000 corresponding to 0.81 Å calibrated pixel size. The movies were collected in 40 frames in defocus range between −0.8 and −2.2 μm with a total dose of 50.19 e^−^/Å^2^ (1^st^ dataset) and 41.19 e^−^/Å^2^ (2^nd^ dataset). Data collection was fully automated using SerialEM^41^.

### Cryo-EM data processing

All data processing operations were performed with RELION-3.0.7^42^ unless otherwise specified.

In total, 17,290 movies of the AVP-V2R-Gs-Nb35 sample at 0.75 mg/ml were collected. Dose-fractionated image stacks were subjected to beam-induced motion correction and dose-weighting using Motioncorr own implementation. Gctf was used to determine the contrast transfer function parameters^43^ from non-dose weighted images. After sorting, micrographs with maximum estimated resolution beyond 5 Å were discarded. Particle picking was carried out using Gautomatch (K. Zhang, MRC LMB (www.mrc-lmb.cam.ac.uk/kzhang/)) allowing to pick out 2,291,432 particles. Particles were extracted in a box-size of 240 Å, downscaled to 4 Å/pix and subjected to reference-free 2D classifications to discard false-positive particles or particles categorized in poorly defined classes. A subset of 1,109,475 particles was selected for further processing. This particle set was subjected to a 3D classification with four classes using the 30 Å low-pass filtered calcitonin receptor map as reference^35^. Particles from the 2 classes representing 27% of total particles and showing a complete AVP-V2R-Gs-Nb35 complex were selected, re-extracted with a pixel size of 1.62 Å, and subjected to a 3D refinement. This subset of 307,125 particles yielded a map with a global resolution (FSC = 0.143) of 4.8 Å resolution. Particles were then subjected to a focused 3D classification without angular and translational alignments with a mask including the complex minus GαsAH. The best class corresponding to 150,000 particles were re-extracted without binning and submitted to a 3D refinement, allowing us to obtain a map at 4.4 Å resolution. All further processing including signal substraction, using of different type of masks, CTF refinement and polishing did not improved the resolution of the map.

In total, 8,490 movies of the AVP-V2R-Gs-Nb35 sample at 4.0 mg/ml were recorded. The image processing steps were the same as previously described, except that the picking was performed using boxnet from Warp software package^44^ allowing us to extract 1,214,575 particles. After a 2D classification to clean the dataset, a subset of 917,990 particles was subjected to two successive rounds of 3D classification. A subset of 150,000 particles was used for further 3D refinements yielding a final map at 4.4 Å resolution.

Both cleaned datasets were merged, corresponding to 1,109,475 particles from dataset 1 and 917,990 particles from dataset 2. Particles were subjected to 3D classification with 3 classes. One class displayed the expected structural features of the AVP-V2R-Gs-Nb35 complex corresponding to 877,003 particles and were selected for a new round of 3D classification with 6 classes. This classification revealed structural variability in the ligand location and at the interface between the receptor and the Gs protein. Three subsets of particles were selected (L, T1 and T2 states), re-extracted with a pixel size of 1.62 Å or 0.81 Å, and subjected to 3D refinements yielding maps at 4.5 Å, 4.7 Å and 5.5 Å respectively. New rounds of 3D refinements were performed in applying a mask to exclude both the micelle and the GαsAH yielding maps at 4.23 Å, 4.4 Å and 4.7 Å. CTF refinement and polishing steps were applied on the three subsets of particles, allowing us to improve the resolution of the best map to 4.17 Å (FSC = 0.143). The 1.62 Å/pixel T1 map was resampled at 0.81 Å/pixel for visualization purposes. Final refinements were processed with the option masking individual particles with zero turned-off. All our attempts to refine our final subsets in cisTEM^45^ and cryoSPARC^46^ using non-uniform refinement did not improve the resolution of final maps.

To investigate the conformational dynamics of the signalling complex, multi-body refinement was performed on 877,003 particles, with two bodies corresponding to AVP-V2R and Gs-Nb35. Local resolution was estimated with the Bsoft 2.0.3 package^47,48^. Map sharpening was re-evaluated with Phenix autosharpen tool^49^. Phenix resolve_cryoEM tool^16^ was used to improve the map interpretability and allowed to increase the estimated resolution to 4.04, 4.13 and 4.5 Å for L, T1 and T2 states, respectively (**Supplementary Fig. 3**).

### Model building and refinement

#### Receptor and AVP initial models

The V2R was built by comparative modelling, using the MODELLER software^50^ and the X-ray structure of the δ-opioid receptor at 3.4 Å resolution (PDB:4EJ4) as a template^51^, sharing a sequence similarity of about 44% with the V2R (on the modeled region). Because modelling loops or terminal regions is a very challenging task and their dynamical behavior is very poorly described in CG simulations, N- and C-terminus of the receptor (residues 1-35 and 335-371, respectively) as well as part of the icl3 loop (237-262) were lacking in the used template. Thus, only residues 36 to 236 and 263 to 334 were modeled. 500 models were generated and the one sharing the best objective function score was further selected as a starting point for the simulations. The disulfide bridge conserved among the class A GPCRs was included between residues 112 and 192 of the V2R.

The AVP peptide (NH3^+^-CYFQNCPRG-CONH2) was built from its X-ray structure available in the PDB (code 1JK4, 2.3 Å resolution) which describes the 6-residues cycle of the peptide in interaction with neurophysin^52^. As depicted in **Supplementary Fig. 4** this structure shows a cycle conformation equivalent to that found in bound (PDB code: 1NPO) and unbound related peptide oxytocin (PDB code: 1XY2)^53, 54^. It was thus preferred to the one describing the trypsin:vasopressin complex (PDB code: 1YF4)^55^ harboring a completely different conformation of the cycle. The three last residues of the peptide (7-PRG-9) were also built with the oxytocin structure templates.

The obtained initial models of both receptor and peptide were then converted to a CG representation using the MARTINI forces field (version 2.2, Elnedyn)^56^. Using such a model, residues (backbone beads) closer than 9.0 Å are bound by a spring displaying a force constant of 500 kJ.mol^−1^.nm^−2^ (default value from the Elnedyn force field). Such a link is meant to maintain both the secondary and the tertiary structures of the polypeptides. The resulting elastic networks of both the peptide and the receptor are represented in **Supplementary Fig. 4**. For the peptide, only the springs involving two residues of the cycle were conserved for further calculations, the three last residues being free to move. The standard elastic network of the receptor was not modified and allowed the latter to open or close freely as no spring was bridging the extracellular loops.

#### MD simulations

The receptor was inserted in a 100×100 Å^2^ lipid bilayer exclusively composed of POPC. To avoid the exploration by the peptide of the intracellular side of the membrane during MD (because of periodic boundary conditions), the system was duplicated/rotated along the z-axis (the two extracellular sides of the receptors were facing each other), in order to create an extracellular compartment. Two copies of the peptide were added to increase the interaction sampling with a 1:1 ratio. In a last step, water and chloride counter-ions were added to neutralize the system. The fully-solvated system included 20004 beads. A representation of the simulated box with two receptors and two peptides is reported in Supplementary Fig. 3. After 10 000 steps of energy minimization using the conjugate gradient algorithm, the system was further equilibrated at 51 different temperatures (in the range 300:450 K by steps of 3 K) in the NVT ensemble, using an integration step of 20 fs and over a period of 5 ns. The final production step was performed in the NPT ensemble, using an integration step of 20 fs and was stopped after 20 μs. During production, Replica-Exchange MD (REMD) was used to improve the sampling of all possible configurations of the peptide:receptor complex. The potential energy difference of adjacent replicas was computed every 1000 steps (20 ps) and their coordinates were exchanged according to a Boltzmann criterion. With the used parameters, the probability of exchange between adjacent replica was in the range 0.11 (300K):0.23(450K). Three independent CG-REMD simulations were run to verify the convergence of the obtained models, together representing a cumulated sampling time of ~3 ms (**Supplementary Fig. 5**). For each of these simulations, a clustering was performed on all conformations of the peptide:receptor complex obtained at the lowest temperature (300 K). To do so, we first concatenated the data corresponding to the four possible complexes (peptide1:receptor1, peptide1:receptor2, peptide2:receptor1 and peptide2:receptor2). For that step, only the conformations displaying at least one peptide:receptor contact were kept (a contact was defined using a cut-off distance of 7 Å). For clustering, we used the algorithm of Daura *et al.*^57^ with a RMSD cut-off of 3.0 Å. The RMSD was computed only on the backbone beads of the peptide’s residues 1 to 6 after structural fit onto those of the V2R. The two cysteines side-chains beads were also included for RMSD calculations. All simulations and analyzes were performed with the Gromacs software (version 5)^58^. Figures were produced with VMD^59^.

#### Refinement of the obtained CG-models in the cryo-EM maps

The Correlation-Driven Molecular Dynamics (CDMD) method^60^ was employed to refine the most populated clusters obtained in CG-REMD using the L state cryo-EM map of the AVP-V2-Gs-Nb35 complex. The principle of the method is to use an accurate force field and thermodynamic sampling to improve the real-space correlation between the modeled structure and the cryo-EM maps. Before this refinement step, the Gs heterotrimer and the Nb35 were modeled using the structure of the β2AR-Gs-Nb35 complex^40^ as a reference. The MARTINI force field restrained the internal conformations of the different partners with an internal elastic network. To increase significantly the conformational plasticity of the receptor and explore new conformations specific to the V2R, we modified its default elastic network. We automatically deleted the “long-range” springs involving two beads whose indexes differ by at least 15. This contributed to delete all inter-helices springs. The resulting elastic network used at this step for the receptor is reported in **Supplementary Fig. 4**. The standard elastic network was conserved for all other partners including the AVP peptide, the G protein and the Nb35. No inter-chains springs were included for the G protein. After conversion of Gs and Nb35 into the CG model, the two proteins were placed at a rational position in respect to the V2R using the β2AR-Gs-Nb35 complex^40^. The full system was inserted in a larger membrane (150×150 Å^2^) and solvated on each side for further calculations.

The fit in each cryo-EM maps was performed in four successive steps summarized in **Supplementary Fig. 6**. First, a quick energy minimization (2000 steps of conjugate gradient) was performed on the full system without taking the map into account. This step was dedicated to the removal of bad contacts resulting from the addition of Gs and Nb35 proteins. Then the second step consisted in a first equilibration of 5 ns (10 fs time-step; NVT; 300 K) performed with CDMD and using a constant targeted low resolution of 5 Å together with a strength constant of 10000 kJ.mol^−1^ for the map potential. This bias was applied only to the backbone beads of the system. This step was useful to quickly optimize the alignment of the system with the targeted map. During this second step, an additional force of 50000 kJ.mol^−1^.nm^−2^ was added to keep the distance between the two centers of masses (COMs) of both the peptide and surrounding residues of the receptor close to its initial value. This force prevented a quick motion of the AVP peptide in the first steps of the simulation that resulted from large forces applying to the receptor. For the subsequent steps of the fitting procedure, this additional force on COMs was removed. During the step 3 (30 ns), the same MD parameters were used but with a gradual increase of both the resolution (from 5 to 3 Å) and the strength constant (from 10000 to 50000 KJ.mol^−1^), over a period of 25 ns. During the last 5 ns these values were kept constant. This step was the key step allowing the whole system to adapt and fit to the maps. Finally, the last step (10 ns) consisted in keeping the resolution and the strength constant at their reached values (3 Å; 50000 KJ.mol^−1^), but this time applying the force only to the backbone and side-chain beads of the peptide. All the other backbone beads of the system were restrained in positions during this step with a force constant of 5000 KJ.mol^−1^. This step was useful to refine the position of the peptide in the density, especially of its side-chains. For every step of the fitting procedure, the fit of each cluster was performed 5 times to verify the convergence of the obtained models (**Supplementary Fig. 7**).

#### All-atom refinement of the models in the maps

The CG-models obtained from the fitting procedure were back-mapped to a full-atom representation. We used the standard “initram” procedure provided by the developers of Martini^61^ with subtle changes. These changes concerned restrains on omega angles and Cα-positions for all chains (V2R, Gs, Nb-35), to keep omega angles in trans conformation and avoid large backbone motions which inevitably would lead to models out of cryo-EM maps. Those restrains were added during the minimization and the MD simulations inherent to the default “initram” procedure. In practice, the “initram” procedure was as follow: 1/ after the very raw guess of atomic positions, from CG-beads, done by the “initram” script, 2/ the Charmm36 force field^62^ was used for 10000 steps of steepest descent disabling the non-bonded terms, 3/ followed by 5000 steps of steepest descent including all terms of the force field, and finally 4/ 300 steps of MD were performed. Except the number of steps, the parameters for minimization and MD simulations were set as default from the “initram” procedure. Minimization and MD simulations were performed using the Gromacs package^58^.

As a final step, iterative manual adjustments were carried out in COOT^63^, and real space refinement using Phenix programs^64^. The model statistics were validated using Molprobity^65^.

### Classical all-atom molecular dynamics simulations

The L state cryoEM structure was subjected to molecular dynamics simulations (MDS). The procedure is described in Supplementary Methods.

### NMR data analysis

The purified V2R was prepared either in neutral amphipol (NAPol)^66, 67^ or in LMNG detergent. In both cases, the V2R was expressed in Sf9 insect cells and purified as described above, except it was cleaved overnight at 4°C using the HRV3C protease at a 1:20 weight ratio (HRV3C:V2R) before concentration and purification by SEC.

1D STD NMR spectra^68^ were recorded either on a mixture of AVP with V2R (400 μM:2 μM), or on AVP (**Supplementary Fig. 8**). Selective methyl resonance saturation was achieved by equally spaced 60 ms Gaussian 180° pulses separated by 1 ms delay at 0 ppm (−50 ppm for reference spectra) at 274K and 283K. An irradiation test was performed on a free peptide sample (400 μM) to verify that only V2R resonances were irradiated. Subtraction of free induction decay with on- and off-resonance protein saturation was achieved by phase cycling. A relaxation delay of 2.6s (Aq+D1) and 128 dummy scans were employed to reduce subtraction artifacts. Investigation of the time dependence of the saturation transfer from 0.5 to 4 s with equally spaced 50-ms Gaussian shaped pulses (separated by a 1-ms delay) showed that 2 s was needed for efficient transfer of saturation from V2R to the AVP. A T1ρ filter of 30 ms was applied to eliminate background resonances of V2R. The transient number was typically 4k. To determine the specificity of STD signals, similar samples were prepared with the antagonist TVP as competitor, using 3 μM V2R, 80 μM AVP and 550 μM TVP. The STD effect was then calculated as (I_0_-I_sat_)/I_0_ where I_0_ and I_sat_ are the intensities of one signal in the reference NMR spectrum and in the on-resonance spectrum, respectively.

We discriminated the different molecular models issued from CG-MD simulations by comparing the experimental STD values and the expected simulated STD from model structures. Back calculation of STD intensities were calculated with the 3.8 version of CORCEMA-ST software^69^. An order parameter value of 0.85 for methyl groups, and a Kon value of a 10^8^ s^−1^ value were used. The correlation times were set to 0.5 and 40 ns for the free and bound states respectively. Calculations with different correlation time values exploring the 0.2-2 ns and 10-30 ns for the free and bound forms, respectively, showed that the simulated profiles, and in particular the correlation coefficient between calculated and experimental values, were much more dependent on the template model than on the correlation time values. Coefficient correlations between simulated and experimental values were calculated for the whole peptide (residues 1-9). Mean correlations factors R1-9 were calculated for five representative structures of each cluster.

### TR-FRET binding assays

V2R binding studies using TagLite® assays (Cisbio, Codolet, France) based on time-resolved-FRET measurements were previously described^70,71^. These are detailed in Supplementary Methods.

### cAMP accumulation assays

As for V2R binding studies, V2R functional studies based on time-resolved FRET measurements were described previously^39, 26^. These are detailed in Supplementary Methods.

### Data Availability

The cryo-EM density maps for the AVP-V2R-Gs-Nb35 complex have been deposited in the Electron Microscopy Data Bank (EMDB) under accession code EMD-12128 (L state) and EMD-12129 (T state). The coordinates for the models of AVP-V2R-Gs-Nb35 complex have been deposited in the PDB under accession numbers 7BB6 and 7BB7 (L and T states, respectively). We declare that the data supporting findings for this study are all available upon request to the corresponding authors.

## Acknowledgements

This work was supported by grants from FRM (grant DEQ20150331736) and ANR (grant ANR-19-CE11-0014) and core funding from CNRS, INSERM and University of Montpellier. The CBS is a member of the French Infrastructure for Integrated Structural Biology (FRISBI) supported by ANR (ANR-10-INBS-05). J.B. was supported by a doctoral fellowship from the Ministère de L’Enseignement Supérieur, de la Recherche et de l’Innovation. We thank the cryo-EM staff at EMBL of Heidelberg (Germany) and the IGF Arpege platform of Pharmacology. We thank Dr. Robert Healey for critical reading of the manuscript and Dr. Amin Sagar for helping in software installation.

## Author contributions

J.B. purified V2R and AVP-V2R-Gs-Nb35 complexes, screened samples by negative staining EM and cryo-EM, prepared grids, collected and processed cryo-EM data, generated the cryo-EM maps and built some Extended data Figures. H.O. managed the Sf9 cell culture and baculoviral infections, expressed and purified V2R, purified AVP-V2R-Gs-Nb35 complexes, prepared grids for cryo-EM. N.F. developed the CG-REMD modelling approach, fitted the models onto the cryo-EM maps and back-mapped the models to all-atom representation. C.L. built the final models of AVP-V2R-Gs-Nb35 complexes into cryo-EM and performed MD simulations. J. L-K-H and A. A. screened samples by negative staining EM and prepared grids for cryo-EM. G.G. contributed to the expression and purification of V2R. J. S-P expressed and purified Gs protein and Nb35 nanobody. S.T. contributed to processing cryo-EM data for generating cryo-EM maps. M.L. contributed to CG-REMD molecular dynamics modelling. S.G. established all the initial procedures for the ternary complex purification. R.S. designed Gs constructs, expressed and purified Gs protein and built most of the Figures. H.D. managed STD NMR experiments and generated the STD NMR data. B.M. designed the V2R construct and determined its pharmacological properties with the help of H.O. S.G. P.B. and B.M. wrote the paper with the input from J.B., N.F. and H.D. Finally, S.G., P.B and B.M. supervised the project.

## Additional information

### Supplementary information

is available in the online version of the paper.

### Correspondence and requests for materials

should be addressed to S.G, P.B or B.M.

---

**Extended data Figure 1.**
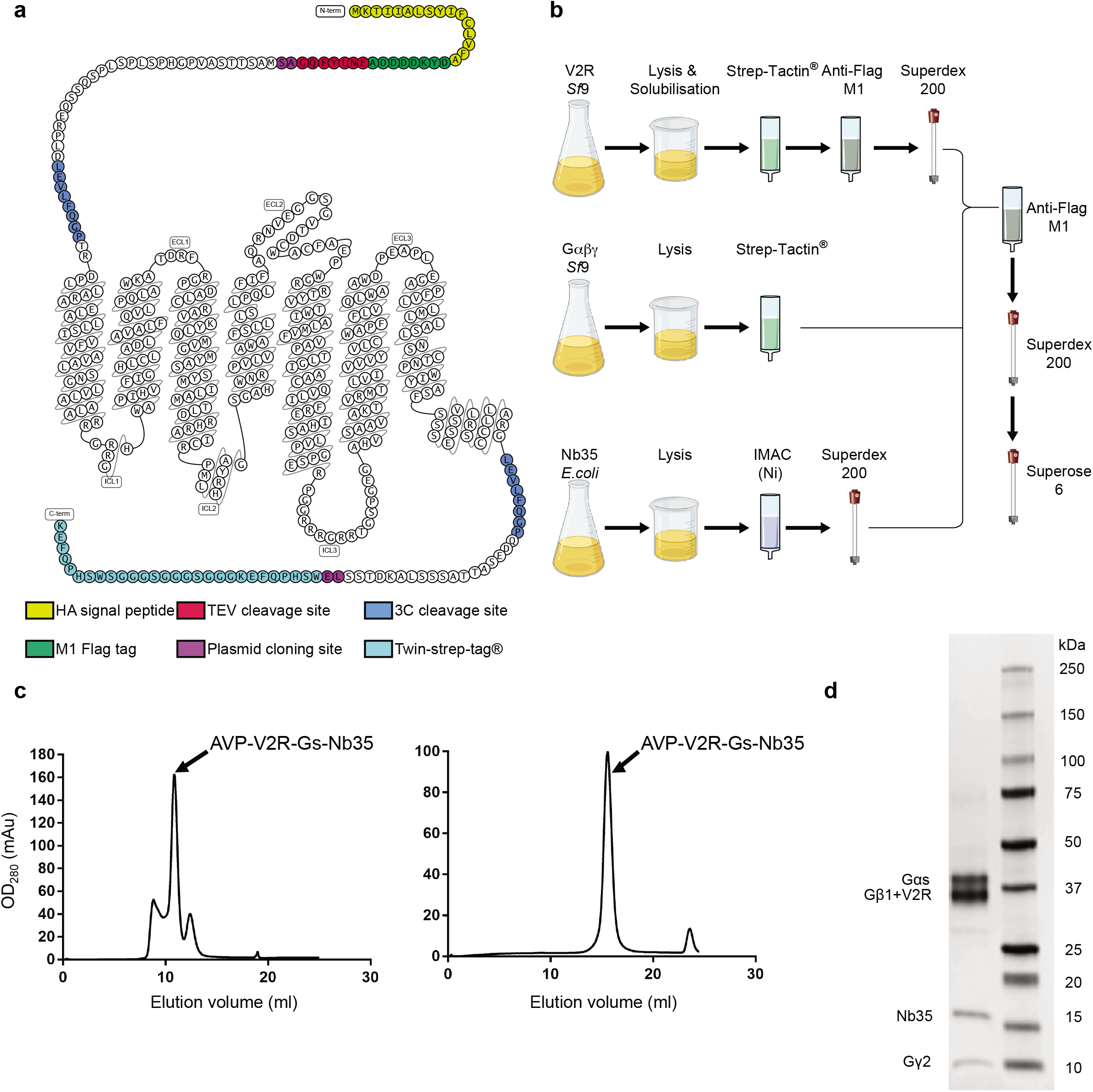
Overview of AVP-V2R-Gs-Nb35 complex preparation and purification. **a**) Modified snake plot from https://gpcrdb.org of the engineered V2R construct used for cryo-EM structure determination. HA, hemagglutinin signal peptide; TEV protease, tobacco etch virus protease; 3C, human rhinovirus 3C protease; plasmid cloning sites are Nhe1 and Xho1 restriction sites. **b**) Workflow for AVP-V2R-Gs-Nb35 assembly. V2R receptor, Gs heterotrimer and Nb35 were expressed and purified separately, the complex being isolated by a final affinity chromatography and two successive SEC steps. **c**) Representative chromatograms of the AVP-V2R-Gs-Nb35 complex using Superdex200 and Superose6 SEC columns show a monodisperse peak. Fractions containing the sample were combined and concentrated for preparation of cryo-EM grids. **d**) SDS-PAGE of peak fraction from the Superose6 step. Coomassie blue staining of proteins confirmed that the complex is made of Gαs, V2R, Gβ1, Nb35 and Gγ2 (AVP is not visible).

**Extended data Figure 2.**
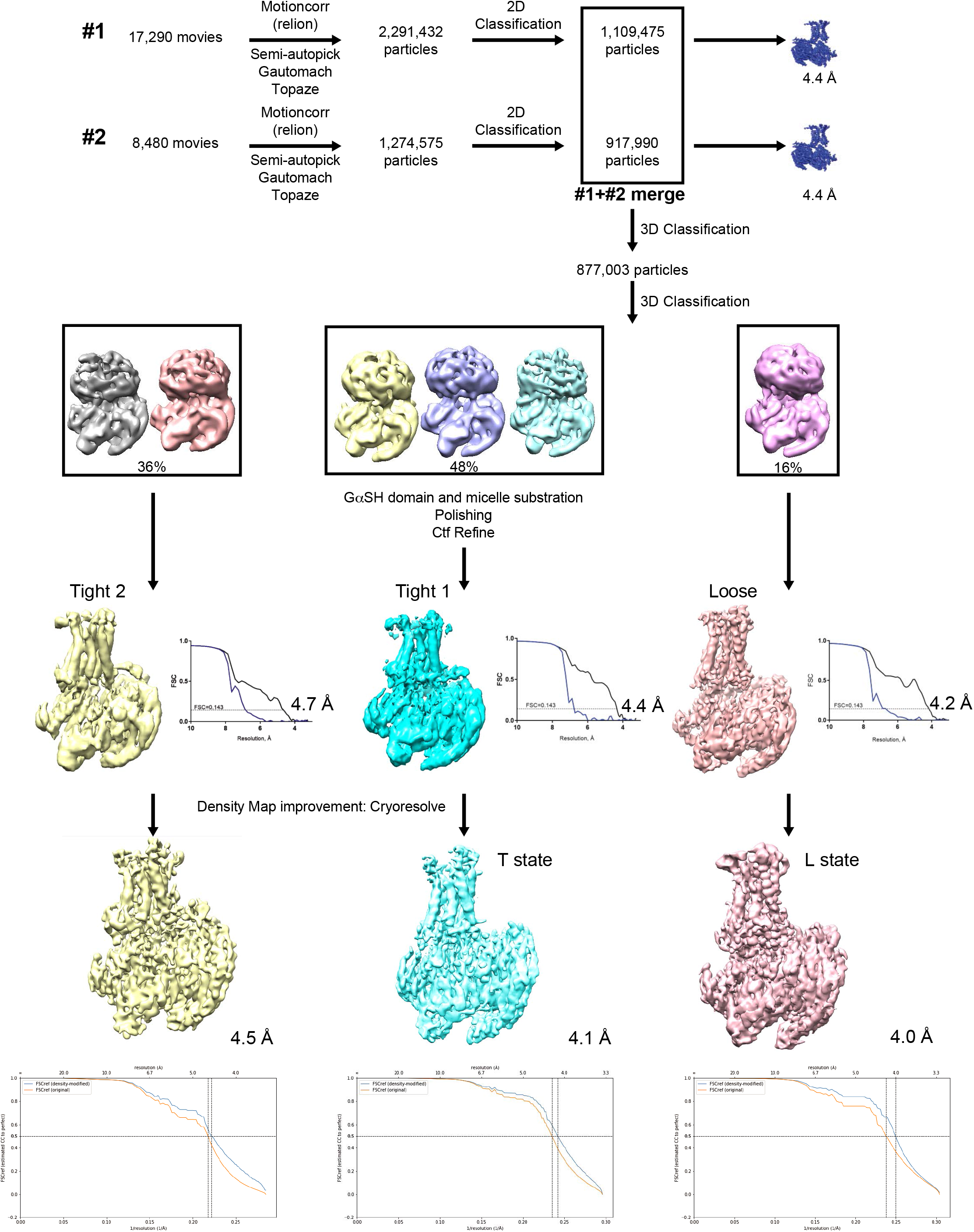
Cryo-EM workflow. Flowchart of the single particle analysis for the two datasets processed separately. Merging of the two sets, substates determination and high-resolution reconstructions. Density map improvement with cryoresolve as a final step.

**Extended data Figure 3.**
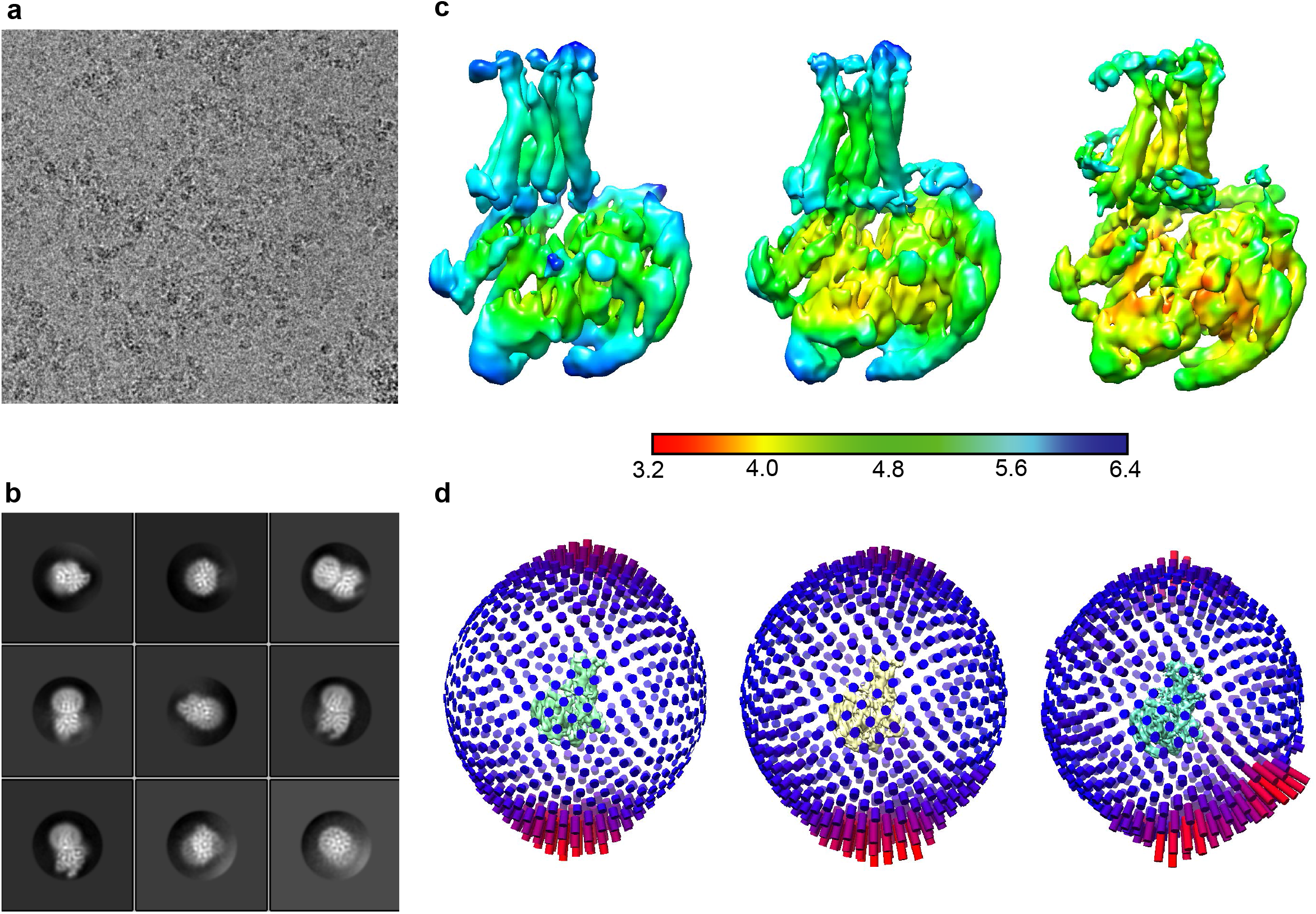
Cryo-EM characterization of the AVP-V2R-Gs-Nb35 complex. **a**) Representative micrograph of the AVP-V2R-Gs-Nb35 protein complex (scale bar, 30 nm). **b**) Representative 2D class averages showing distinct secondary structure features (including the V2R TM regions embedded in the detergent micelle) and different orientations of the AVP-V2R-Gs-Nb35 complex (scale bar, 5 nm). **c**) Local resolution estimation computed with blocres from bsoft program; Tight-2, Tight-1 and Loose particle density maps are shown, respectively. **d**) Euler angle distribution of particles from the final reconstructions for Tight-2, Tight-1 and Loose populations.

**Extended data Figure 4.**
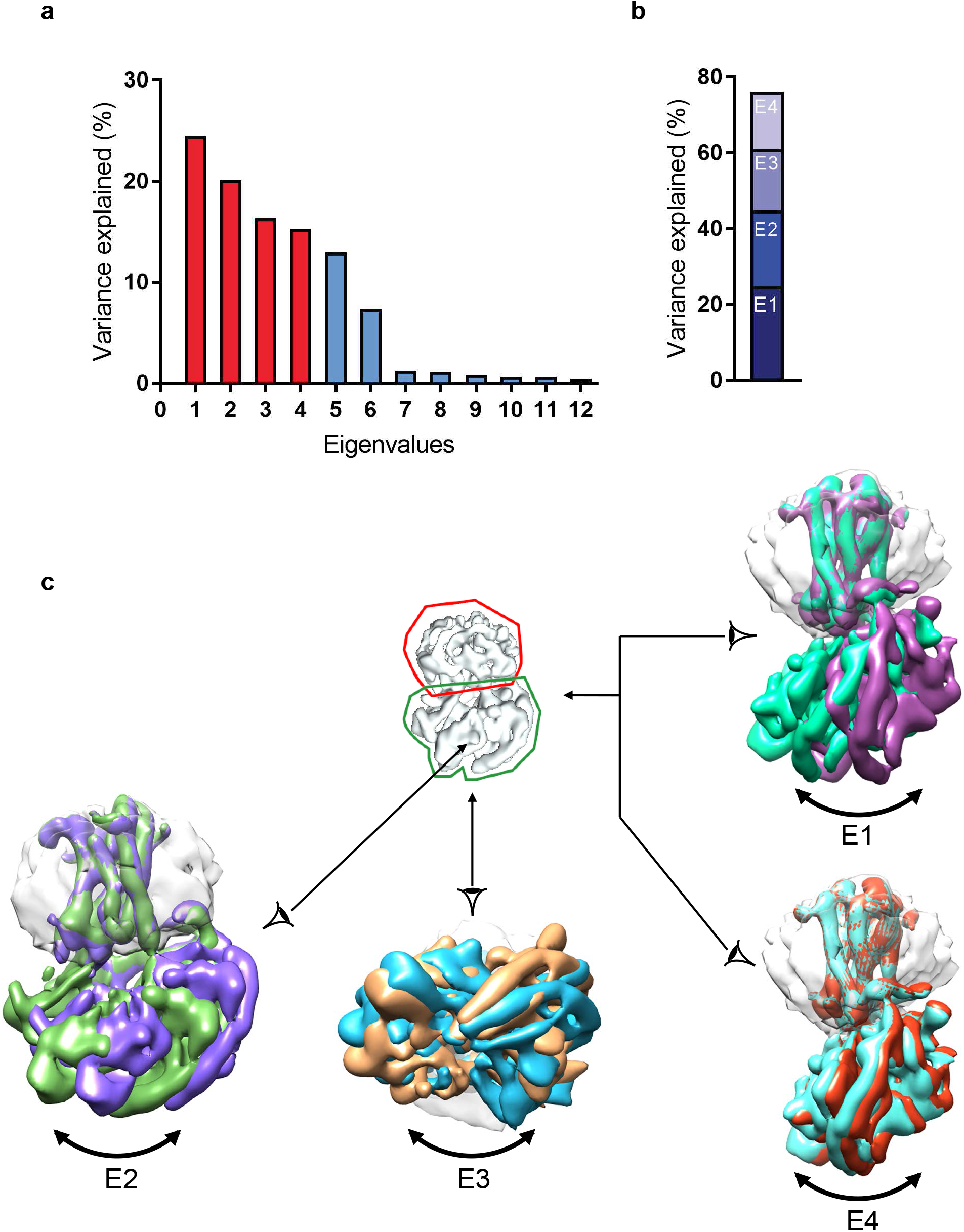
Flexibility in the AVP-V2R-Gs-Nb35 complex. **a**) The contribution of each of the twelve eigen-vectors (numbered along x-axis) to the variance of the overall final map is illustrated. **b**) Eigenvectors 1 to 4 correspond to 78% of the variance. **c**) Mask employed for multi-body refinement is shown in the middle, detergent micelle and V2R are surrounded by a red line, Gs and Nb35 by a green line. Maps corresponding to the four first vectors are illustrated, showing swing-like motion and tilting of Gs-Nb35 with respect to AVP-V2R. E4 is part of E1.

**Extended data Figure 5.**
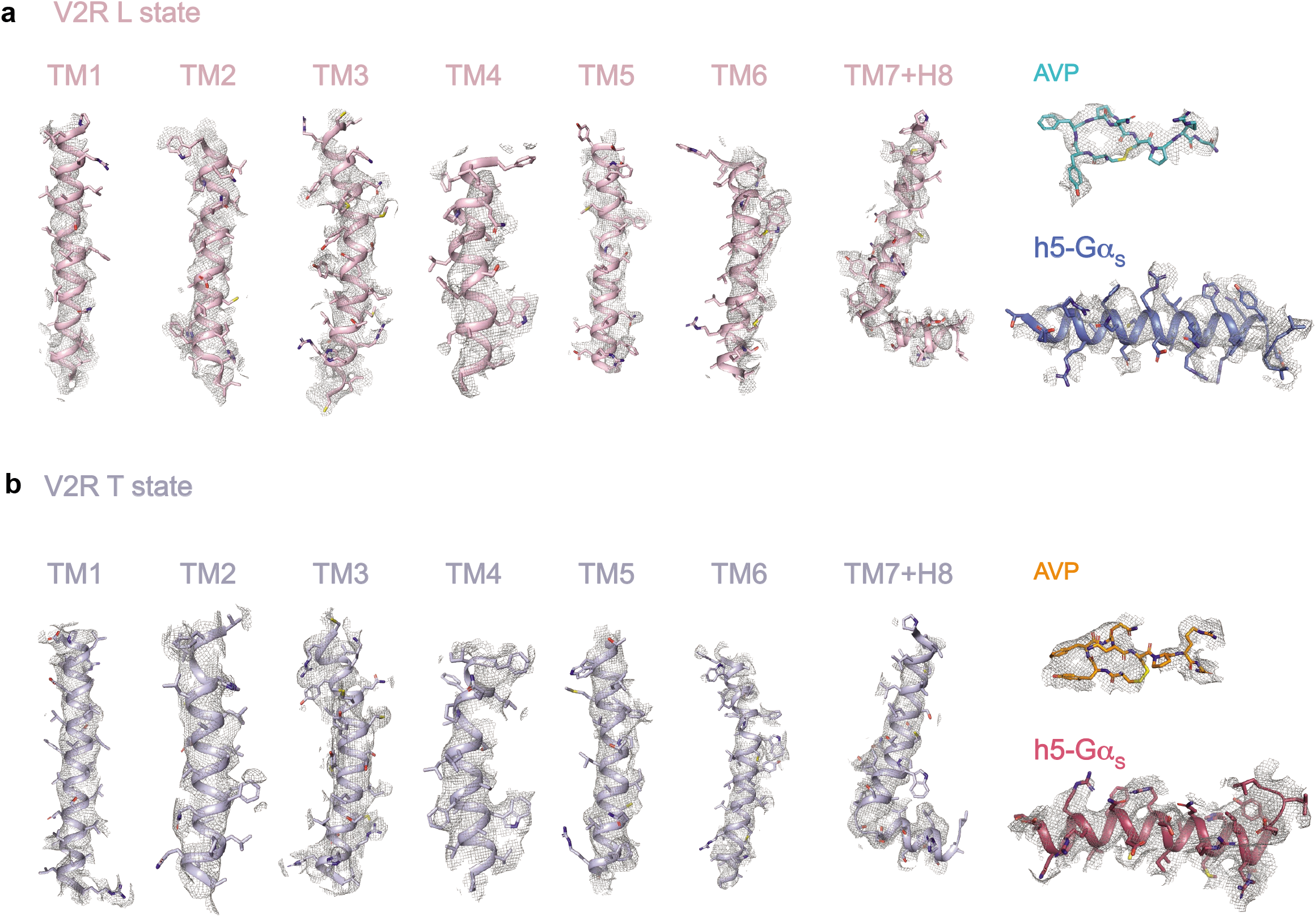
Cryo-EM map quality for L and T states of the AVP-V2R-Gs-Nb35 complex. **a**) The density and model for TM helices 1-7 and helix H8 of V2R, α5 helix of Gαs and AVP in Loose (L) state. **b**) The density and model of corresponding protein domains and AVP in the Tight (T) state.

**Extended data Figure 6.**
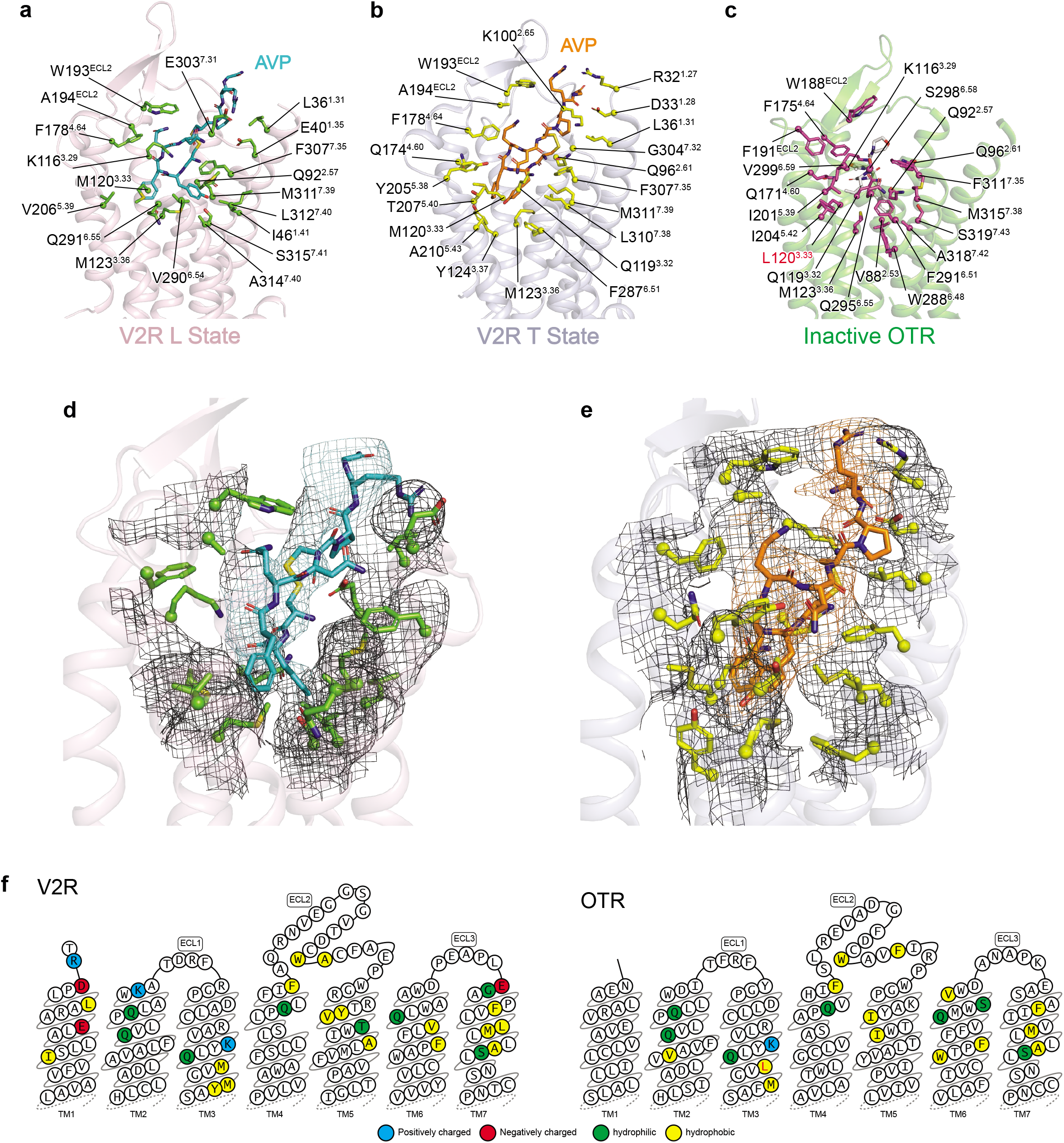
V2R and OTR binding prockets: binding of AVP vs Retosiban. AVP binding poses are viewed from the side of V2R helix bundle in L (**a**), T (**b**) states and are compared with that of retosiban (in white sticks) in OTR (**c**). Receptor residues directly interacting with the ligands (at a maximum 5 Å distance) are indicated (Ballesteros-Weinstein numbering). In the OTR, L120^3.33^ (highlighted in red) is a mutation introduced in the sequence to increase thermostability and facilitate crystallogenesis (V120L). Densities of V2R residues in contact with AVP are shown in L (**d**) and T states (**e**), respectively. **f**) Residues of V2R and OTR involved in the binding of ligands are shown in receptor snake-like plot representations (https://gpcrdb.org).

**Extended data Figure 7.**
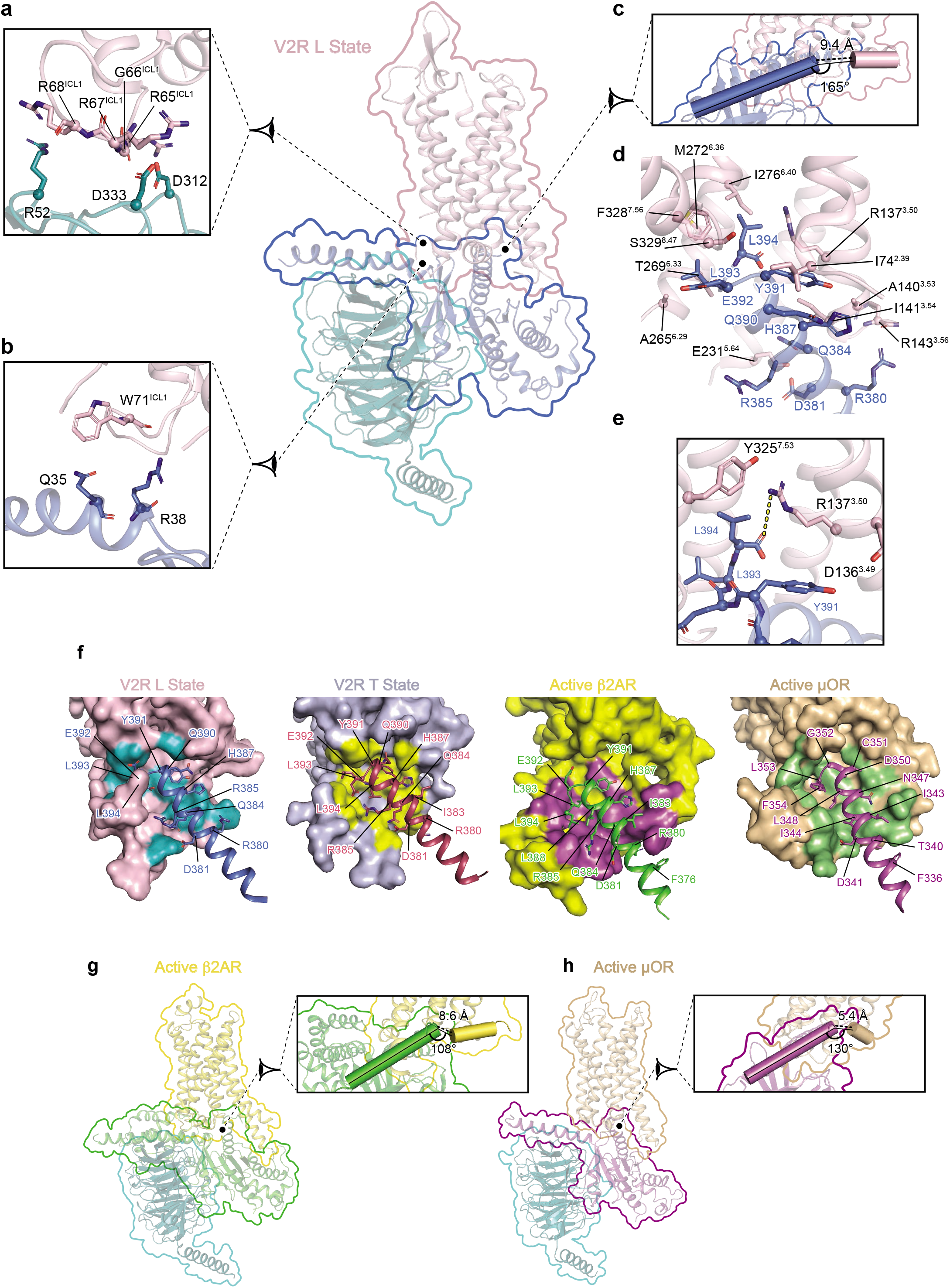
Interface of the V2R L state with Gs. Interactions between V2R and Gs heterotrimer are shown. Specific interfaces are depicted, and residues in close proximity (within a maximal 3.9 Ä distance) are highlighted (pink for V2R, turquoise for Gβ subunit, blue for Gα subunit). **a**) Interaction of V2R icl1 with Gβ subunit. **b**) Interaction of V2R icl1 with N-terminal helix of Gα subunit. **c**) Positioning of C-terminal h5 helix of Gα subunit relative to V2R helix 8. The distance between Gα and helix 8 is indicated. Angle between these two domains is shown. **d**) Interacting residues between the C-terminal h5 helix of Gα subunit and V2R. **e**) Zoom on an ionic bridge between the C-terminal free carboxylic moiety of h5 helix of Gα subunit and V2R R137^3, 50^. **f**) Comparison of class A GPCR-Gα protein interfaces. The V2R-Gαs interfaces of L and T states are compared to those of the β2AR-Gαs and μOR-Gαi complexes. The h5 helix of the Gα subunit is shown for each complex with its residues indicated. The residues of receptors in contact with the Gα C-terminal are coloured. **g**) and **h**) Position of the C-terminal h5 helix of Gα subunit relative to receptor helix 8 in active β2AR-Gs and μOR-Gi complexes, respectively. Distances and angles between these domains are indicated as in panel c.

**Extended Data Movie 1 | Representation of the flexibility of the signalling complex.** This animation represents the first four eigenvectors computed by multi-body refinement analysis, which explain 78% of the variability of the cryo-EM projections (see Extended Data Fig.4).

**Extended Data Movie 2 | PCA analysis obtained from molecular dynamics simulations.** This animation shows the motion described by the first 3 eigenvectors from a PCA analysis starting from the L model. The PCA was performed on protein Cα atoms using MD trajectories representing about 2.6 μs of aggregated simulation time (see Supplementary Fig. 9).

**Extended data Table 1:**
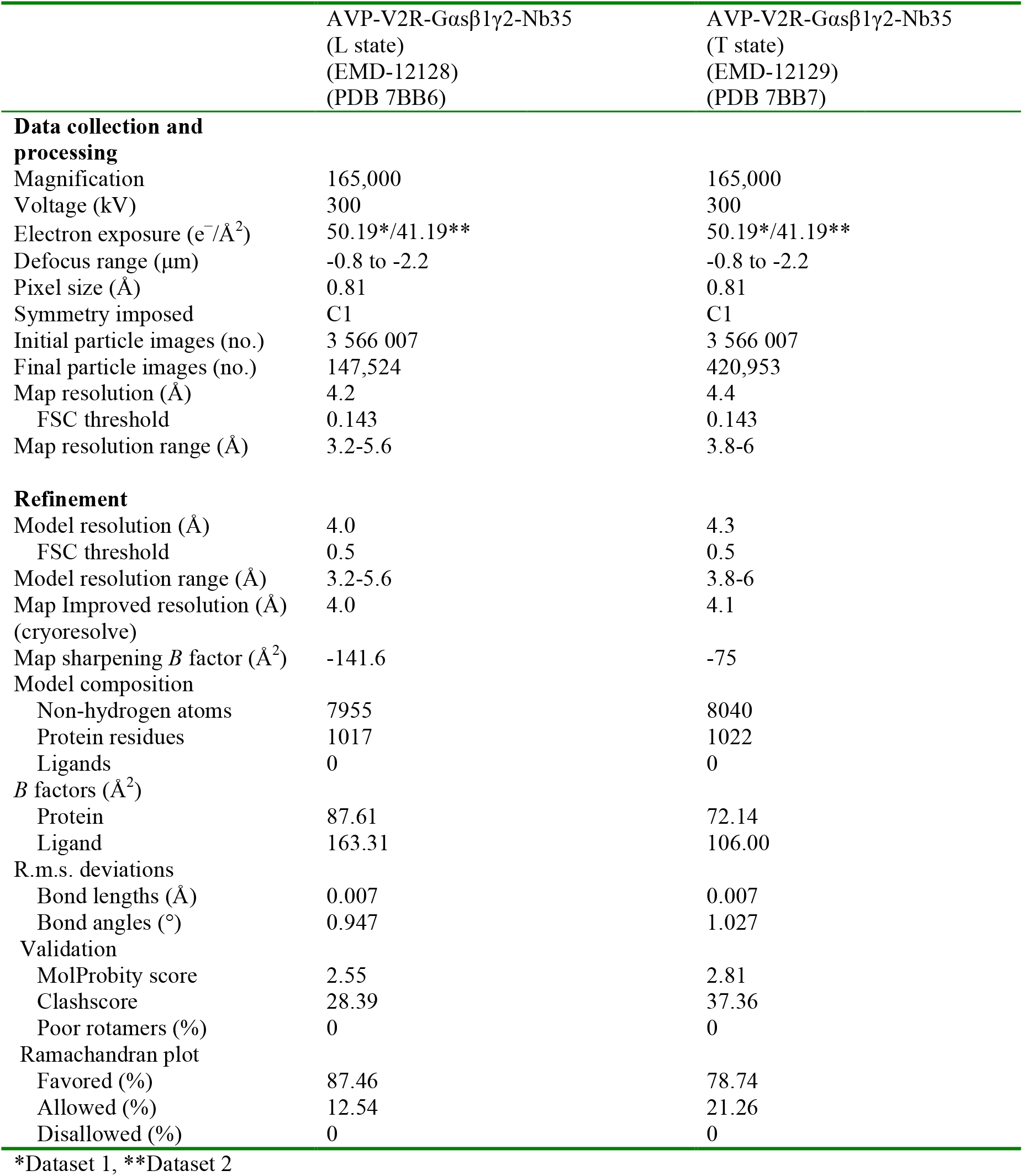
Cryo-EM data collection, refinement and validation statistics

