## Supplementary material for "Structure of the antidiuretic hormone vasopressin receptor signalling complex": SI

### 1/ Supplementary Methods

#### V2R construction

The optimized sequence of the human V2R was cloned into a pFastBac™1 vector (Invitrogen) for insect cell expression. To facilitate expression and purification of the V2R construct used for cryo-EM (**Extended data Fig. 1**), the hemagglutinin signal peptide (MKTIIALSYIFCLVFA) followed by a Flag tag (DYKDDDDA) were added at the N-terminus, and a Twin-strep-tag® (WSHPQFEKGGGSGGGSGGGSWSHPQFEK) inserted at the C-terminus. In addition, N22 was substituted with a glutamine residue to avoid N-glycosylation, and C358 mutated into an alanine to eliminate potential intermolecular disulfide bridges during solubilization and purification. A TEV protease cleavage site (following the Flag tag) and 2 HRV3C protease cleavage sites (1 inserted in the N-terminus between D30 and T31), the other inserted in the C-terminus between G345 and Q354 and replacing R346-TPPSLG-P353) were also added to remove N- and C-termini and facilitate structure determination. M1L2 residues were replaced by AS residues, and LE residues were added just before the Twin-strep-tag®, during subcloning (introduction of a Nhe1 and a Xho1 restriction site, respectively). Sequence modifications did not affect the receptor ligand binding or function (**Supplementary Fig. 1**).

#### Nb35 expression and purification

The production and purification of Nb35 were performed following a protocol established by Kobilka and coworkers[1]. Nanobody-35 (Nb35) having a C-terminal 6His-tag was expressed in the periplasm of *E. coli* strain BL21 following induction with 1 mM IPTG. Cultures of 2L were grown to $OD_{600}$ to 0.6 at 37°C in LB medium containing 0.1% glucose and 100 µg/ml ampicillin. Induced cultures were grown overnight at 25°C. Cells were harvested by centrifugation and lysed in ice-cold buffer (50 mM Tris-HCl pH 8, 125 mM Sucrose, 2 mM EDTA). Lysate was centrifuged to remove cell debris and Nb35 was purified by nickel affinity chromatography. Eluate was concentrated to 5 mg/ml and loaded onto a Superdex 200 (GE HealthCare, 16/600) at a 1 ml/min flowrate. Fractions containing the monodisperse peak of Nb35 were pooled and dialysed overnight against buffer 10 mM HEPES pH 7.5, 100 mM NaCl at RT°C. The dialysed sample was concentrated to approximately 100 mg/ml using a 10 kDa MWCO concentrator (Millipore). Aliquots were stored at -80°C until use.

#### Negative stain microscopy observations

Three µl of AVP-V2R-Gs-Nb35 complex at 0.04 $mg.mL^{-1}$ were applied 2 min on glow-discharged carbon-coated grids, and then negatively stained with 1% uranyl acetate for 1 min. Observation of EM grids was carried out on a JEOL 2200FS FEG operating at 200 kV under low-dose conditions (total dose of 20 electrons/Å2) in the zero-energy-loss mode with a slit width of 20 eV. Images were recorded on a 4K X 4K slow-scan CCD camera (Gatan Inc.) at a nominal magnification of 50,000X with defocus ranging from 0.5 to 1.5 µm. Magnifications were calibrated from cryo-images of tobacco mosaic viruses. In total, 37 Micrographs were recorded allowing us to pick 22,791 particles using e2boxer from Eman2 package[2] (**Supplementary Fig. 2a**). Further processing was done with Relion 2.0[3,4]. The particles were subjected to a 2D classification included to get rid of free micelles and dissociated components of the complex (**Supplementary Fig. 2b**). From 2D classes, 14,545 particles corresponding to the V2R-Gs-Nb35 complexes were selected, representing 63% of all particles. This selection was used to calculate an ab-initio low-resolution model (**Supplementary Fig. 2c**). The sample was also subjected to NS-EM analysis after 5 days. At this point after particles picking and 2D classification, 35% of particles were representing the complex. The fresh sample was also mixed with 100µM GTPγS and 10 µM of SR121463 V2R antagonist, and visualized in negative stain to observe complete dissociation (**Supplementary Fig. 2d**).

**Classical all-atom molecular dynamics simulations**

The system was set up using the CHARMM-GUI Micelle builder[5]. The protein complex (L state) was inserted into a hydrated, equilibrated micelle composed of 60 molecules of Lauryl Maltose Neopentyl Glycol after addition of missing protein loops in Coot. 495 sodium and 511 chloride ions were added to neutralize the system, reaching a final concentration of approximately 150 mM. MDS were performed in GROMACS 2020 using the CHARMM36m force field and the CHARMM TIP3P water model. The input systems were subjected to energy minimization, equilibration and production simulation using the GROMACS input scripts generated by CHARMM-GUI[6]. Briefly, the system was energy minimized using 5000 steps of steepest descent, followed by 375 ps of equilibration. NVT (constant particle number, volume and temperature) and NPT (constant particle number, pressure and temperature) equilibrations were followed by NPT production runs. The van der Waals interactions were smoothly switched off at 10–12 Å by a force-switching function[7], whereas the long-range electrostatic interactions were calculated using the particle mesh Ewald method[8]. The temperature and pressure were held at 310.15 K and 1 bar, respectively. The assembled system was equilibrated by the well-established protocol in Micelle Builder, in which various restraints were applied to the protein, detergents and water molecules, and the restraint forces were gradually reduced during this process. During production simulations an NPT ensemble was used with isotropic pressure coupling via the Parrinello-Rahman barostat method while the Nose-Hoover thermostat was used to maintain a temperature of 310.15 K. A leapfrog integration scheme was used, all bonds were constrained and hydrogen mass repartitioning was applied[9], allowing for a time step of 4 ps to be used during NPT equilibration and production MDS. We performed 10 independent production runs starting from the highest resolution L state model, for a total simulation time of ~ 2.6 µs. Production runs were subsequently pooled together, and the resulting trajectory was analyzed using GROMACS tools to yield principal components. The analysis was performed on the subset of Cα atoms common to the simulated and experimental structures using 1 frame per nanosecond of trajectory. The experimental L, T1 and T2 states were included in the analysis for comparison.

**TR-FRET binding assays**

V2R binding studies using TagLite® assays (Cisbio, Codolet, France) based on time-resolved-FRET measurements were previously described[10,11]. Briefly, HEK cells were plated in white-walled, flat-bottom, 96-well plates (Greiner CELLSTAR plate, SigmaAldrich) in DMEM containing 10% Fetal Calf serum (Lonza), 1% non-essential amino acids and penicillin/streptomycin (GIBCO) at 15,000 cells per well. Cells were transfected 24h later with a plasmid coding for the V2R version used in cryo-EM studies fused at its N-terminus to the SNAP-Tag (SNAP-V2R) (Cisbio Bioassays, Codolet, France). Transfections were performed with Xtreme-GENE 360 (SigmaAldrich), according to the manufacturer recommendations: 10 µL of a premix containing DMEM medium (qsp 10 µl/well), Xtreme-GENE 360 (0.3 µl/well) (SigmaAldrich), SNAP-V2 coding plasmid (30 ng/well) and non-coding plasmid (70 ng/well) was added to the culture medium. After a 48h culture period, cells were rinsed once with Tag-lite medium (Cisbio Bioassays,

Codolet, France) and incubated in the presence of Tag-lite medium containing 100 nM benzyl-guanine (BG)-Lumi4®-Tb for at least 60 minutes at 37°C. Cells were then washed four times. For saturation studies, cells were incubated for at least 4 hours at 4°C in the presence of Benzazepine-red (BZ) nonpeptide vasopressin antagonist (BZ-DY647, Cisbio Bioassays, Codolet, France) at various concentrations ranging from $1x10^{-10}$M to $1x10^{-7}$M. Nonspecific binding was determined in the presence of 10µM vasopressin. For competition studies, cells were incubated for at least 4 hours at 4°C with benzazepine-red ligand (5 nM) and increasing concentrations of vasopressin ranging from $1x10-^{11}$M to $3.16x10^{-6}$M. Fluorescent signals were measured at 620nm (fluorescence of the donor) and at 665nM (FRET signal) on a Pherastar (BMG LABTECH, Champigny s/Marne, France). Results were expressed as the 665/620 ratio (10000*(665/620)). Specific variation of the FRET ratio was plotted as a function of benzazepine-red concentration (saturation experiments) or competitor concentration (competition experiment) (**Supplementary Fig. 1a,b**). All binding data were analyzed with GraphPad 8.3.0 (GraphPad Software, Inc.) using the one site-specific binding equation. All results are expressed as the mean ± SEM of at least three independent experiments performed in triplicate. $K_i$ values were calculated from $IC_{50}$ values with the Cheng-Prusoff equation.

**cAMP accumulation assays**

As for V2R binding studies, V2R functional studies based on time-resolved FRET measurements were described previously[12,13]. Briefly, CHO cells were plated in six-well plates (Falcon) at 350,000 cells per well and transfected 24 h later with JetPei (Ozyme) with a pRK5 plasmid coding for the version of the V2R used in the cryo-EM studies. A mix of isotonic NaCl solution (200 µl/well) containing JetPei (Ozyme, 2 µl / well), V2R coding plasmid (1ng/well) and non-coding plasmid (3000 ng/well) were added to the culture medium (2 ml). 24 h later, cells were harvested with trypsin and cultured in white-walled, flat-bottom, 96-well plates (Greiner CELLSTAR plate; Sigma-Aldrich) at a density of 30,000 cells per well in DMEM containing 10% Fetal Calf serum (Lonza), 1% non-essential amino acids and penicillin/streptomycin (GIBCO). After a 24 h culture period, cells were treated for 30 minutes at 37°C in the cAMP buffer with or without increasing AVP concentrations ($3.16x10^{-12}$ to $10^{-6}$M) in the presence of 0.1 mM RO201724, a phosphodiesterase inhibitor (SigmaAldrich). The accumulated cAMP was quantified using the cAMPdynamic 2 Kit (Cisbio Bioassays - Codolet, France) according to the manufacturer protocol. Fluorescent signals were measured at 620 nm and 665 nm on a Spark® 20M multimode microplate reader (Tecan). Data were plotted as the FRET ratio (10000*(665/620)) as a function of AVP concentration (log[AVP]). Data were analyzed with Graphpad Prism using the "Dose-response – stimulation" subroutine (**Supplementary Fig. 1c**). EC50s were determined using the log(agonist) *vs*. response – variable slope (4 parameters) fit procedure. Experiments were repeated at least three times on different cultures, each condition in triplicate. Data are presented as mean±SEM.

### 2/Supplementary Discussion

To improve the expression of the human V2R and facilitate its purification, we constructed a receptor version with a hemagglutinin signal peptide followed by a flag tag at its N-terminus, and with a twin strep tag at its C-terminus (**Extended data Fig. 1**). In addition, N22 was substituted with a glutamine residue to avoid N-glycosylation, and C358 mutated into an alanine to eliminate possibility of intermolecular disulfide bridges. It is important to note here that, apart from receptor engineering designed uniquely for expression and purification purpose, and, unlike many of the recently published GPCR structures, we did not modify the receptor sequence (the V2R is wild-type from T31 to G345). Our aim was to avoid possible artefacts and irrelevant information due to the introduction of mutations in the transmembrane core domain of the receptor, even if this was at the expense of lower resolution cryo-EM data. Moreover, and before the recombinant expression of the receptor in Sf9 insect cells, the pharmacological properties of the engineered V2R were verified in HEK mammalian cells (**Supplementary Methods**). The cryo-EM version of the V2R bound a fluorescent nonpeptide antagonist as well as AVP with high affinity (**Supplementary Fig. 1**, $K_d$ and $K_i$ = 2.27 ± 0.24 nM (n=3) and 1.12 ± 0.5 nM (n=3), respectively), close to the values determined for a WT V2R[11]. Moreover, the receptor was proven to be functional as it was able to stimulate cAMP accumulation upon AVP binding ($K_{act}$ = 2.05 ± 0.11 nM (n=4), like the WT V2R in transfected cells[14] (**Supplementary Fig. 1**).

Images of the complex first recorded in NS-EM revealed a homogeneous distribution of the particles, as observed from 2D class averages (**Supplementary Fig. 2**). More than 60% of the particles correspond to the complex. A reconstruction at 20 Å clearly showed the micelle of detergent and the G protein-Nb35 components. Fitting the 3D model of the crystal structure of the β2-adrenergic receptor-Gs-Nb35[1] in this low-resolution reconstruction map confirms that V2R-Gs-Nb35 displays typical structural features of a transmembrane signaling GPCR complex (**Supplementary Fig. 2**). Moreover, the addition of the specific V2R nonpeptide antagonist SR121463[15] and GTPγS to the purified complex led to the dissociation of the different components (**Supplementary Fig. 2**), confirming the functionality of the signaling particle. Taking into account these different results, the samples were then vitrified onto cryo-EM grids for structure determination.

To improve the resolution of the different cryo-EM density maps, we used the recent algorithm Cryoresolve[16], based on the maximum-likelihood density modification used to improve maps from macromolecular X-ray crystallography. This analysis allowed us to increase the resolution of L, T1 and T2 density maps from 4.2, 4.5 and 4.7 Å respectively, to 4.0, 4.1 and 4.5 Å. The application of density modification to cryo-EM map L of the AVP-V2R-Gs-Nb35 complex enhanced the visibility of many details for some V2R TM regions, for the hormone AVP, for helix h5 of the Gαs subunit and for Gβ2 subunit (**Supplementary Fig. 3**). In many cases, the improvement of the density map enabled to identify more accurately side chains of amino-acid residues.

To optimize hormone docking in the cryo-EM density map, we combined different complementary approaches, i.e. receptor:AVP computational molecular dynamic simulations and experimental STD NMR. First, the conformational sampling of the peptide:receptor complex was improved using the unbiased CG method coupled to

Replica-Exchange MD (REMD) protocol. We successfully employed this protocol to predict the binding modes of peptides in the class A GPCRs NTS-1, CXCR4 and GHSR[17,18]. The simulated system is described in **Fig. 1** and **Supplementary Fig. 4**. Three independent CG-REMD simulations were run, together representing about 3 ms of cumulated simulation time. Each of the three simulations led to respectively 288, 306 and 302 clusters of peptide:receptor conformations. Interestingly, the first 10 most populated clusters reported in **Fig. 1** were identically retrieved among the three independent simulations as shown by the RMSD matrix reported in **Supplementary Fig. 5,** and represented more than 60% of the whole explored conformations. After addition of the Gs heterotrimer and Nb35 proteins, refinement of each of these clusters was performed in the L cryo-EM density map. At this step, we used the CDMD method[19] while keeping advantage of using a CG representation for sampling speed and better agreement with the resolution of the maps (**Supplementary Fig. 6**). Fitting of each cluster was repeated 5 times. Typical curves of Cross Correlation coefficients as a function of time for each CDMD are reported **in Supplementary Fig. 7.** It shows that the used protocol reached a "plateau" in each case indicating the convergence of the fit for all clusters. Small variability of the position of the peptide among the five obtained models for clusters 2 and 5 (mean RMSD of 3.0 and 2.2 Å, respectively) and in a lower manner for the clusters 6 and 8 (mean RMSD of 3.2 and 3.6 Å, respectively) was seen (**Fig. 1**). The higher values obtained for the other clusters (in the range 4.8 to 8.7 Å) were explained by the upper starting position of the peptide in the pocket, finding more easily the density located at the surface of the receptor during the fitting procedure. Finally, the CG models obtained from the fitting procedure were back-mapped to an all-atom representation. Minimization, MD simulations, iterative manual adjustment and real space refinement were carried-out to finalize AVP docking.

The AVP binding modes were further cross-validated using experimental STD NMR spectroscopy which can efficiently monitor the binding and map the contact surface of a given ligand with its cognate GPCR[20,21]. 1D STD spectra were thus recorded either on a mixture of AVP with V2R or on AVP alone (see **Methods**, **Supplementary Fig. 8**). Intense STD signals were only observed in the presence of V2R, mostly for the aromatic protons of $Tyr^2$ and $Phe^3$ residues of AVP. The addition of the orthosteric antagonist Tolvaptan (TVP) significantly attenuated the STD signals, demonstrating specific binding of AVP to the V2R orthosteric site (**Supplementary Fig. 8**). Calculation of normalized STD effects as $I_{STD}$-$I_{ref)}$/$I_{ref}$ (see **Methods**) showed that the most intense effects were indeed observed for the N-terminal cyclic part of AVP, with a strong involvement of the aromatic side chains of $Tyr^2$ and $Phe^3$ (and to a lesser extend $Cys^1$), whereas the residues in the C-terminal tripeptide ($Pro^7$, $Arg^8$ and $Gly^9$-amide) were less impacted upon V2R binding (**Fig. 1**). In addition, we compared these experimental STD values to the expected STD values from AA models issued from MD simulations and subsequently refined with the density maps. As explained in Methods, coefficient correlations between simulated and experimental STD values were calculated for the whole peptide ($R_{1-9}$). As shown in **Fig. 1**, cluster 5 fitted on L density map appeared as the best cluster fitting to experimental STD values.

### 3/ Supplementary Figures

Supplementary Figure 1: Pharmacological and functional properties of the cryo-EM V2R construct.

**a**) Binding of the benzazepine-red fluorescent antagonist to the V2R construct measured by FRET (see Methods). Specific binding of BZ-Red from a typical saturation assay is shown as FRET ratio (665nm/620 nm). The experiment was repeated 3 times each point measured in triplicate. Each value is presented as mean ± SEM. **b**) Binding of AVP to the V2R construct is illustrated as FRET ratio (665nm/620nm). Specific binding of benzazepine-red is shown. The fluorescent antagonist was used at 5 nM with or without increasing concentrations of AVP. A typical competition curve is shown, and was repeated at least 3 times each point in triplicate. Each value is presented as mean ± SEM. **c**) Capacity of the V2R construct to functionally activate adenylyl cyclase measured by FRET (see Methods). The cAMP accumulation is shown as FRET ratio (665nm/620nm) and measured in the presence of increasing concentrations of AVP. A typical experiment is shown, was repeated at least 3 times, each point in triplicate. Each value is presented as mean ± SEM.

Supplementary Figure 2: Negative stain electron microscopy characterization of the AVP-V2R-Gs-Nb35 complex.

**a**) Representative micrograph of the purified sample of the complex isolated from the Superose6 SEC peak (scale bar, 54 nm) **b**) 2D most representative class averages showing different orientations. (scale bar, 18 nm). **c**) Density map of the AVP-V2R-Gs-Nb35 complex (contour level set to 0.115), and fitting of the 3D model of the crystal structure of β2ARGs-Nb35 complex in this low resolution map. **d**) Representative micrograph of AVP-V2R-Gs-Nb35 complex dissociated using an excess of 10 μM SR121463 (selective nonpeptide antagonist of the V2R) and 100 μM GTPγS (scale bar, 43 nm).

Supplementary Figure 3: Improvement of the L density map using cryoresolve.

In all panels, the density map and the corresponding final all-atom 3D model are superimposed. The improved map (right) is compared to the original map (left). V2R is depicted in purple, AVP in grey, Gαs subunit in orange and Gβ2 subunit in green. Increase in the visibility of several different regions of the AVP-V2R-Gs-Nb35 complex is shown: for instance contacts between F214 and F287 in V2R TM5 and 6 (**a**), W164 in V2R TM4 (**b**), AVP (**c**), N-terminal α helix of Gαs subunit (**d**) and H62-W63 in Gβ2 subunit (**e**).

Supplementary Figure 4: Coarse grain-REMD molecular dynamics approach

**a**) Structural alignment of the two X-ray structures available for AVP (PDB codes: 1JK4 and 1YF4). The 1JK4 structure which describes the 6-residue cycle of AVP was preferred to the 1YF4 structure corresponding to the full length peptide, because it displays a cycle conformation equivalent to tat found in the unbound and bound oxytocin peptide analog (1XY2 and 1NPO). **b**) Schematic representation of the internal elastic networks used for the peptide AVP. **c**) Schematic representation of the internal elastic network used for V2R (side and extracellular view). **d**) The full system used for the CG-REMD simulations included 2 receptors and 2 ligands in order to create an artificial extracellular compartment and improve the conformational sampling of the AVP:V2R complex. **e**)

Modified elastic network of the receptor used for the fit of the obtained CG models into the cryo-EM density maps.

Supplementary Figure 5: CG-REMD simulations
**a**) Cross-RMSD matrix of the ten most populated clusters resuting from the 3 independant CG-REMD simulations and showing that the same models were systematically retrieved (white squares correspond to RMSD < 3Å). **b**) Analyse of the populations of all the obtained clusters in terms of cumulative sums showing that the ten first clusters together represent more than 60% of the whole conformations. Data from the 3 independant simulations are reported in blue, orange and grey, respectively.

Supplementary Figure 6: Summary of the successive steps employing the CDMD method to fit the models resulting from the CG-REMD simulations into the cryo-EM maps.

Supplementary Figure 7: Typical curves of cross-correlation coefficients as a function of time for each CDMD simulation.
**a**) and **b**) Cross-correlation coefficients values computed between the experimental and the simulated maps along CDMD simulations starting from the 10 most observed orientations of AVP in its receptor. In **a**), we reported one representative cross-correlation coefficients curve for each cluster whereas in **b**), the five curves obtained for the same cluster are depicted (five independent replicas). In each case, it shows a small variability of the obtained values and the convergence of the models at the end of the protocol. These curves were extracted from the main step 3 of fitting procedure (see Supplementary Figure 5).

Supplementary Figure 8: Mapping of AVP interaction surfaces by STD NMR experiments.
**a**) and **b**) Comparison of standard 1D proton spectrum (AVP 400 μM) with STD experiments on (**a)** 400 μM AVP and 400 μM AVP binding to 2 μM V2R and (**b)** 80 μM AVP binding to 3 μM V2R in absence/presence of 550 μM TVP (tolvaptan). Buffer resonance (Bis-Tris) and detergent resonance are labelled buf and det, respectively.

Supplementary Figure 9: Principal components analysis (PCA) obtained from molecular dynamics simulations of the L state.
**a)** Collective motions of the AVP-V2R-Gs-Nb35 complex captured by PC1, PC2 and PC3. Motions are illustrated as linear interpolations between the extreme projections of the structures onto the PCs. Each cylinder, therefore, describes the path of each Cα atom between the extremes (on a red–white–blue color scale). **b)** and **c)** Conformational landscape of the AVP-V2R-Gs-Nb35 complex in PC space for the MD ensemble. 2D projections of MD trajectories along PC1 and PC2 (b) or PC1 and PC3 (c) were converted into 2D histograms to represent the density of population of each conformational state of the complex. The experimental L, T1 and T2 states are shown in white as (o), ($x_1$) and ($x_2$), respectively. **d)** Cumulative fraction of variance captured by the first 10 eigenvectors.

Supplementary Figure 10: Activation motifs and coupling interfaces in the AVP-V2R-Gs-Nb35 complex with the details of the cryo-EM maps.
In all panels, the 3D models and the corresponding cryo-EM density maps (shown as green, orange or blue mesh) are superimposed. Close-up views of activation motifs in L

and T states are shown on the left (**a**, **b**, **c**), whereas V2R-Gs contacts in the T state are shown on the right (**d**, **e**, **f**). The toggle switch CWxP motif including $W284^{6.48}$ (**a**), the PSY transmission motif with $P^{5.50}$-$S^{3.40}$-$Y^{6.44}$ residues (**b**) and the conserved NPxxY motif with $Y325^{7.53}$ (**c**) are highlighted. Different interaction interfaces stabilizing the V2R-Gs complex are depicted: the ICL1 of V2R is in close proximity with the Gβ subunit (**d**) and with the N-terminal helix of Gα subunit (**e**) of Gs protein, the unusual ionic bridge contact of V2R $R137^{3.50}$ side-chain (DRH motif) is in close proximity with the free carboxylic acid function of the C-terminal extremity of Gα subunit of Gs (**f**). The L or T conformation of V2R is depicted in pink or blue grey, respectively. The Gβ subunit is shown in turquoise, and the Gα subunit of Gs is illustrated in raspberry.

**Supplementary Figure 1: Pharmacological and functional properties of the cryo-EM V2R construct**

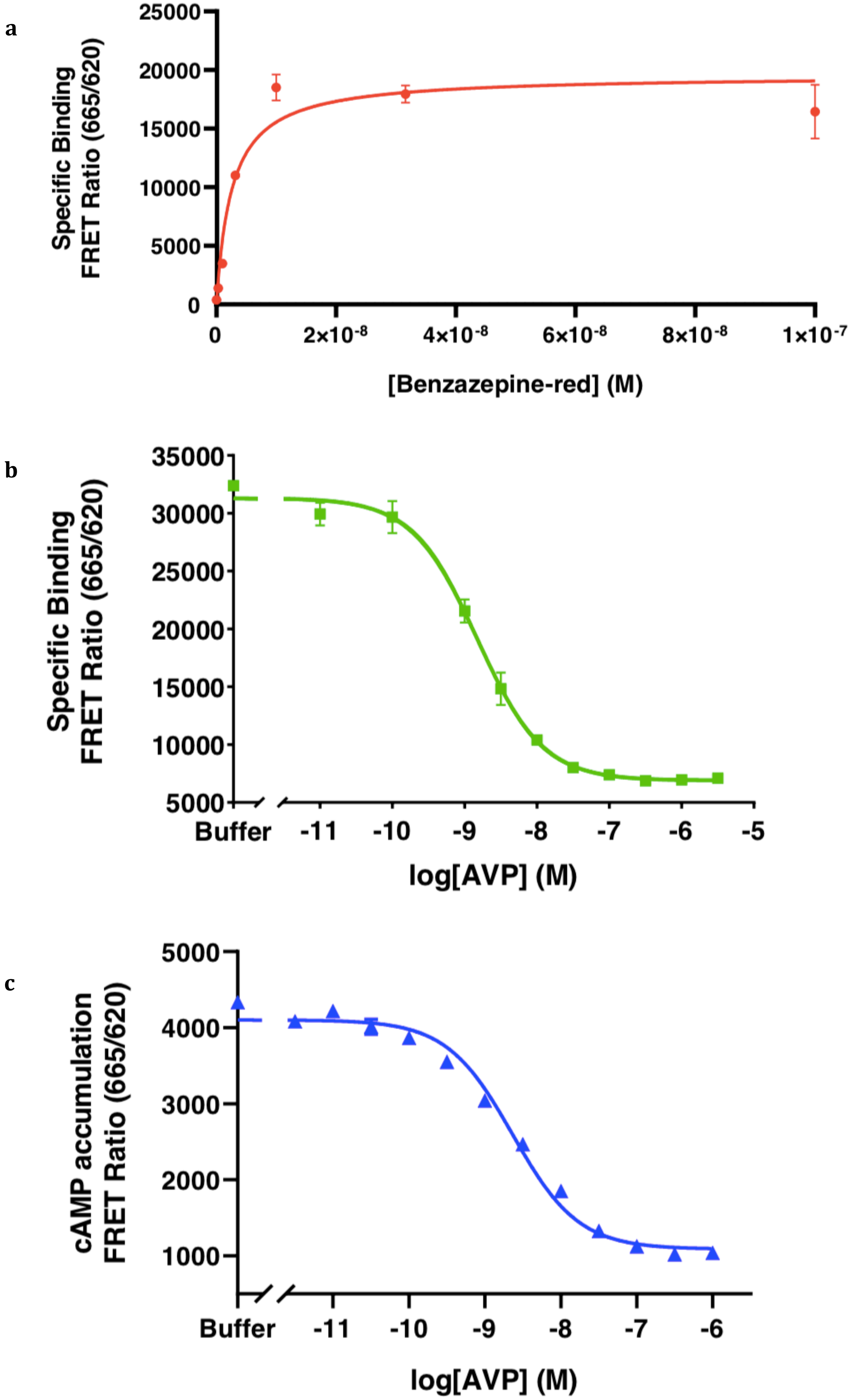

**Supplementary Figure 2: Negative stain electron microscopy characterization of the AVP-V2R-Gs-Nb35 complex**

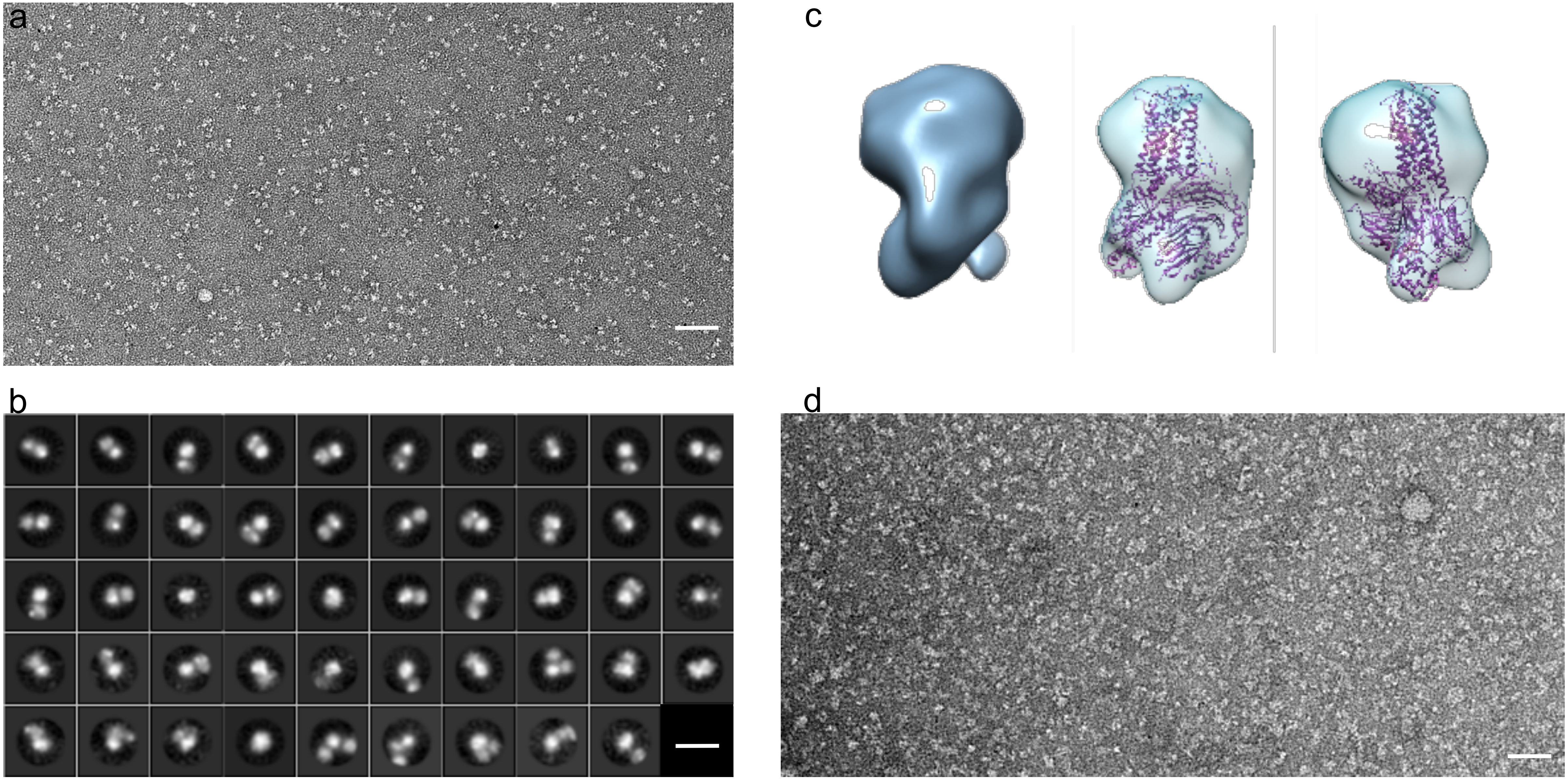

**Supplementary Figure 3: Improvement of the L density map using Cryoresolve**

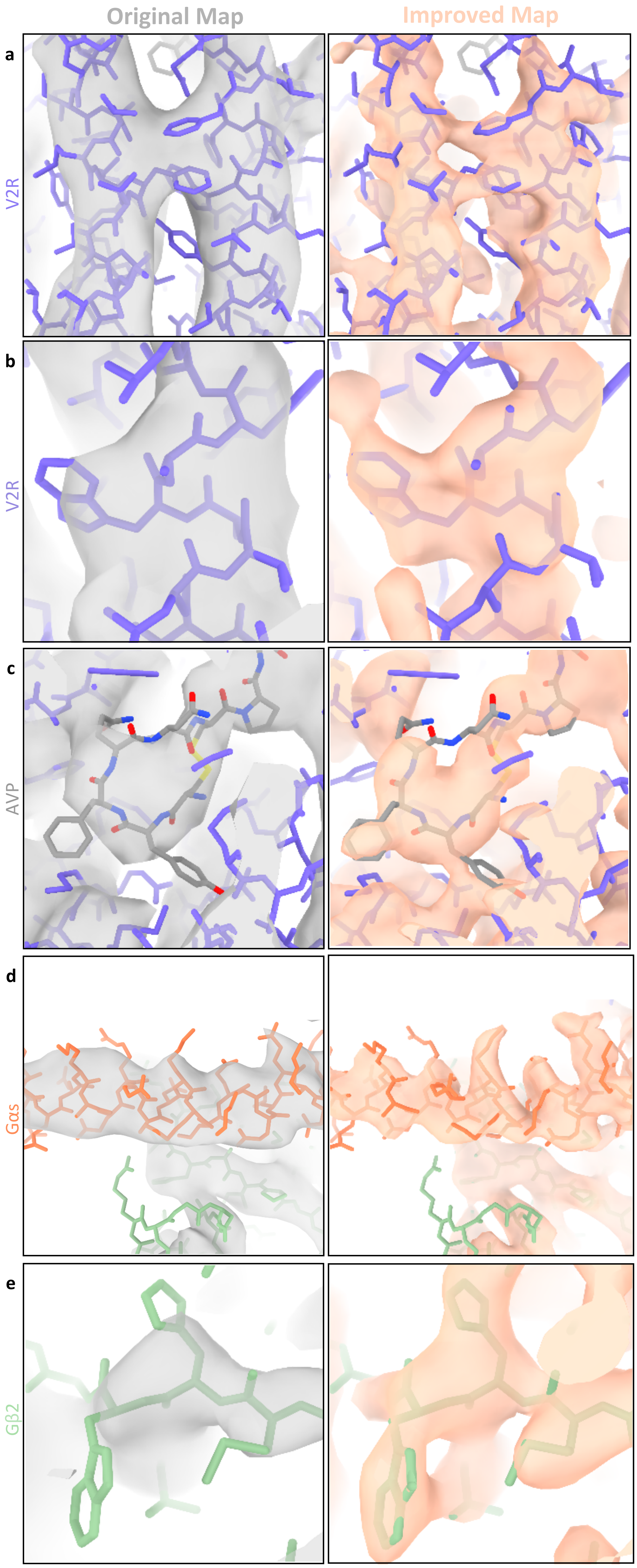

Supplementary Figure 4: Coarse-grain-REMD molecular dynamic approach

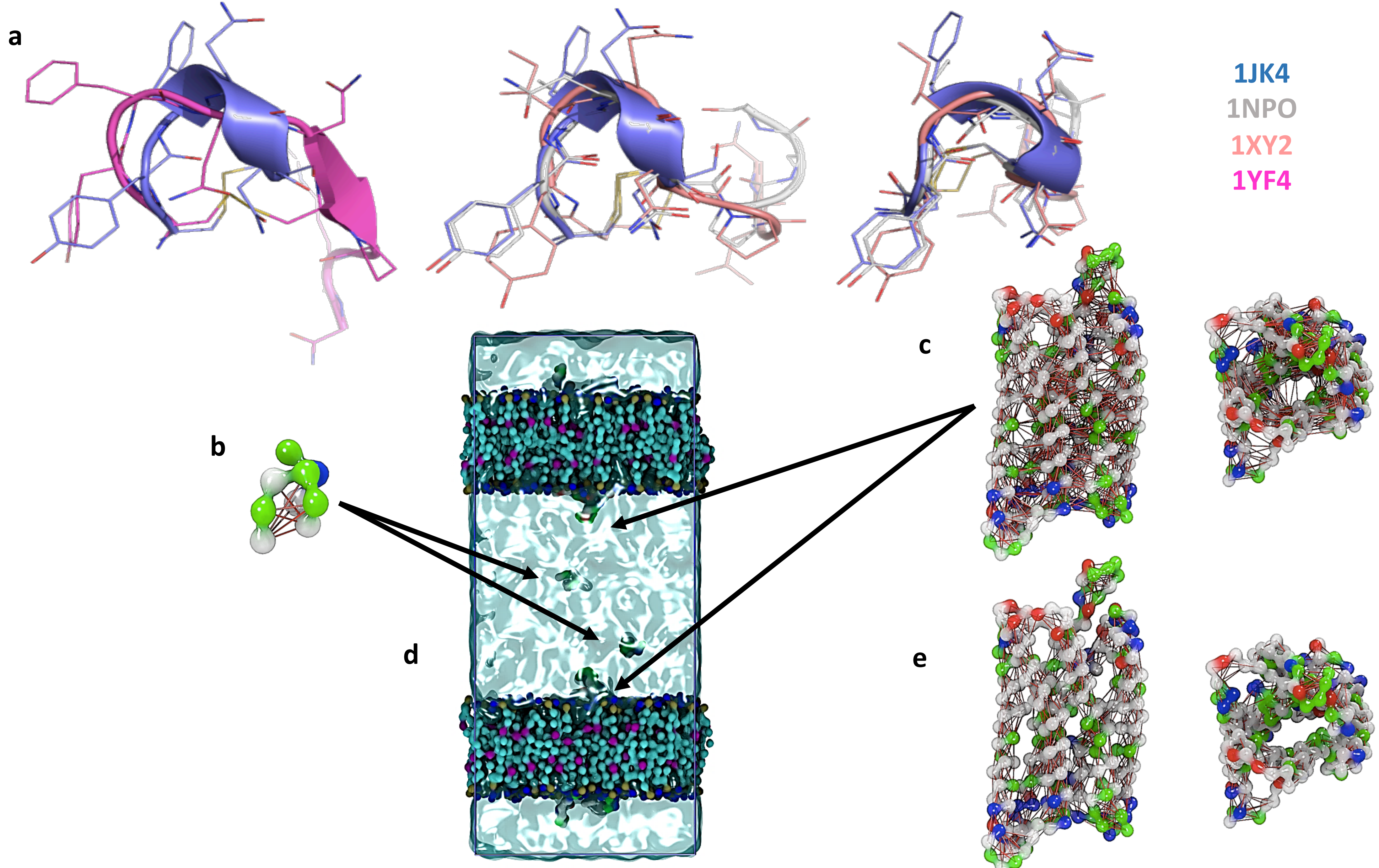

**Supplementary Figure 5: CG-REMD simulations**

a

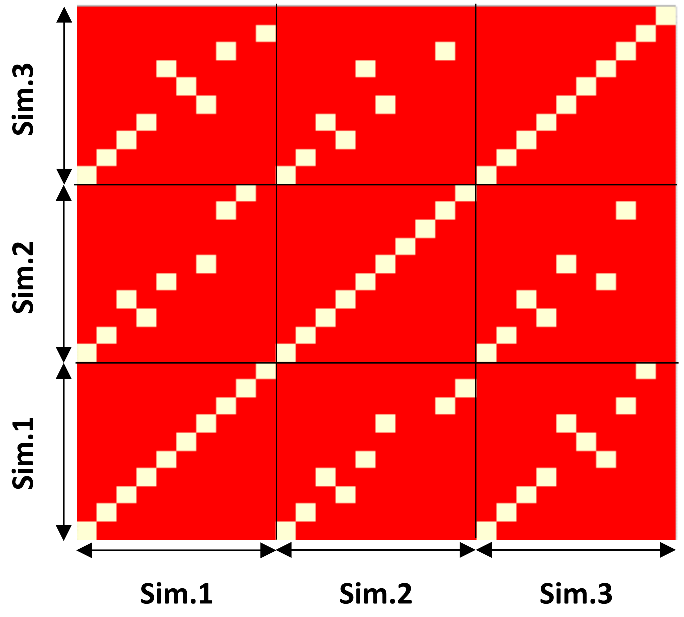


b

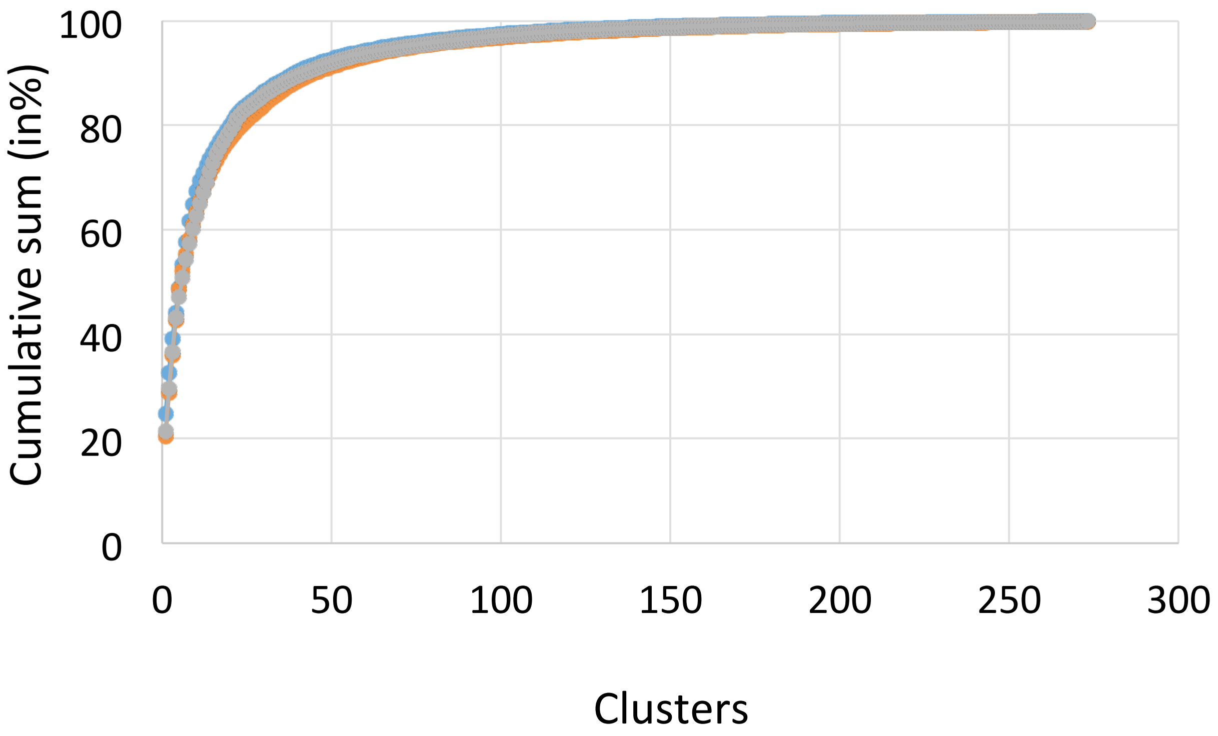


c

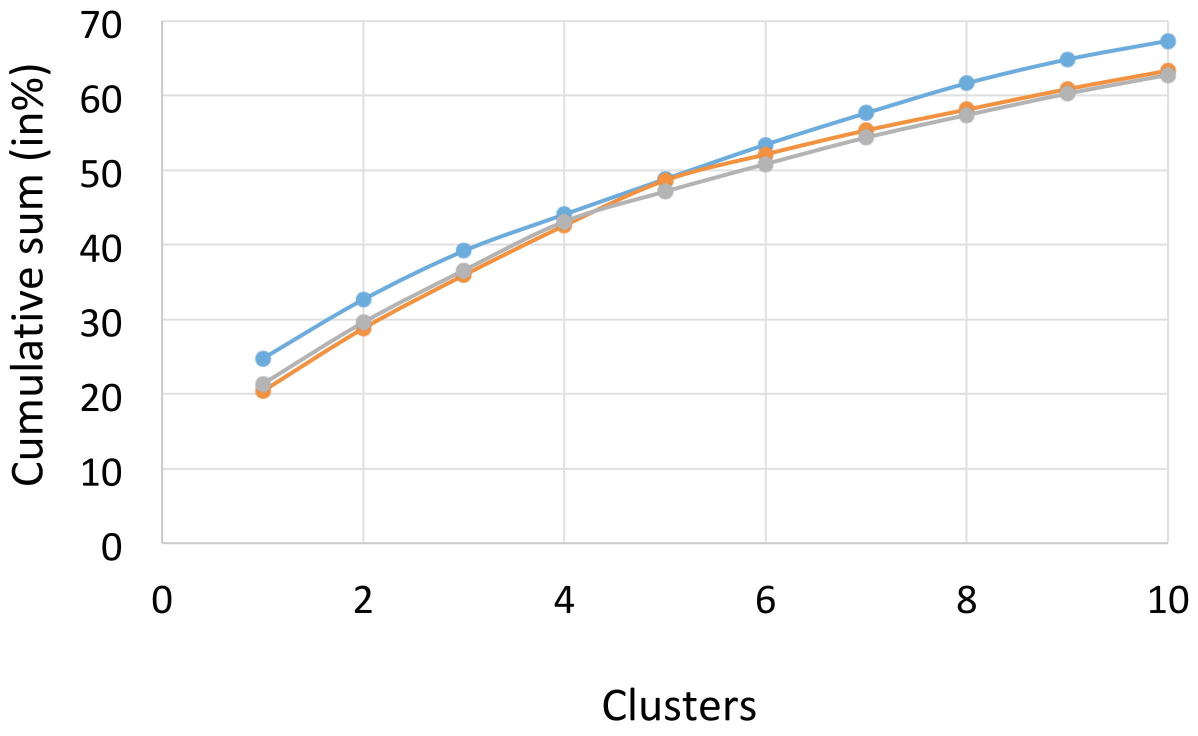

**Supplementary Figure 6: Summary of the CDMD method to fit the models into the cryo-EM maps**

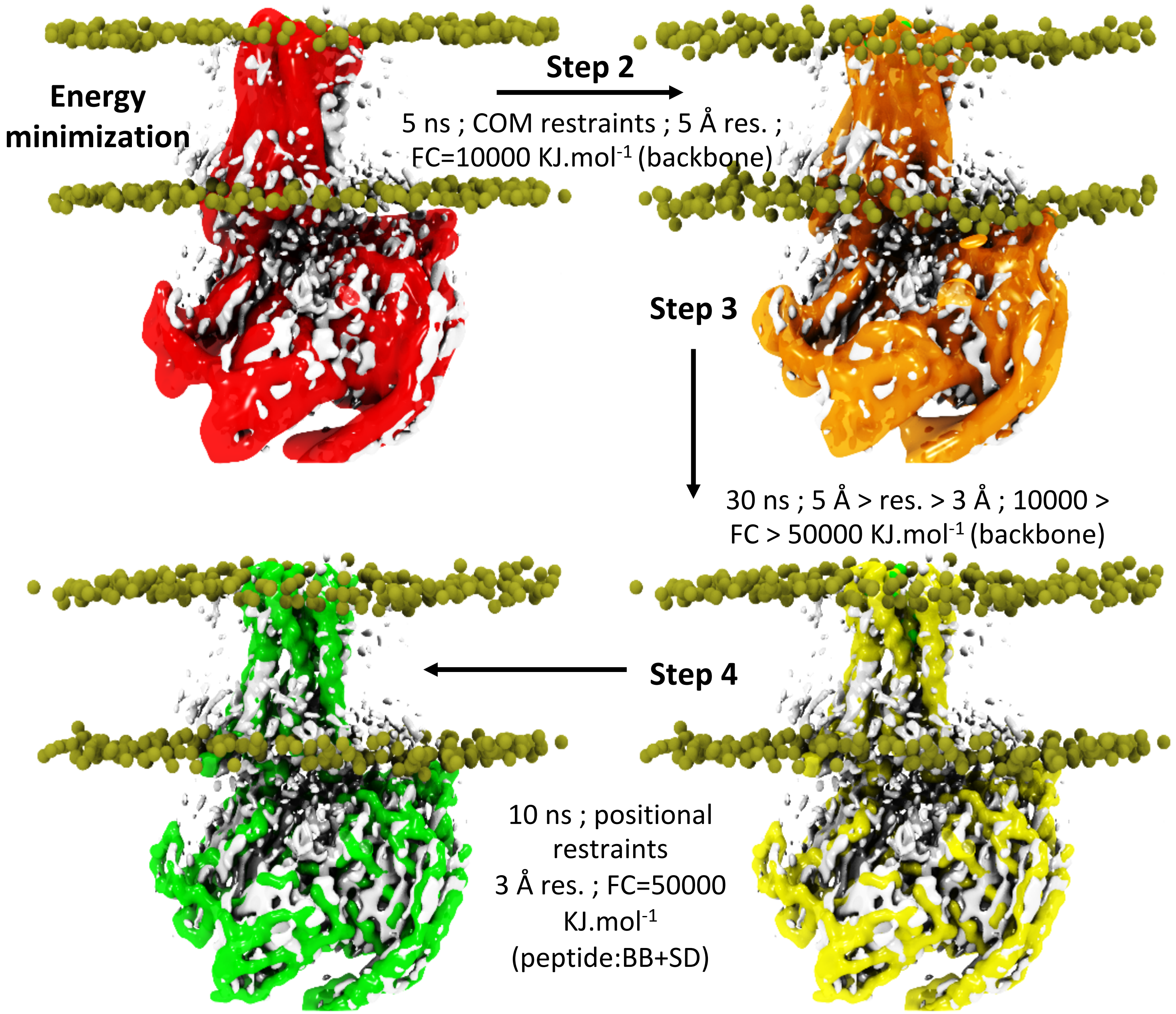

**Supplementary Figure 7: typical curves of cross-correlation coefficients as a function of time for each CDMD simulation**

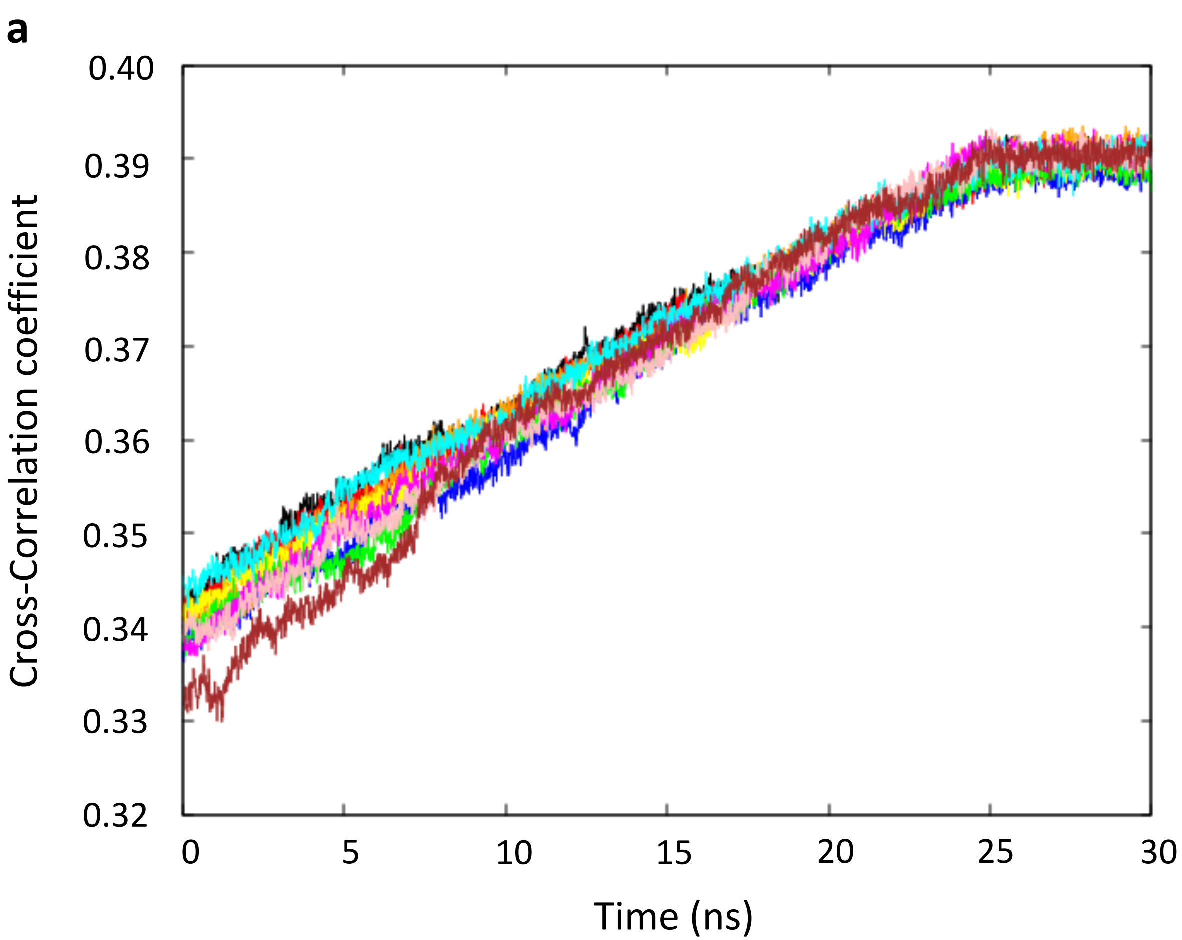

**Supplementary Figure 8: Mapping of AVP interaction surfaces by STD NMR experiments**

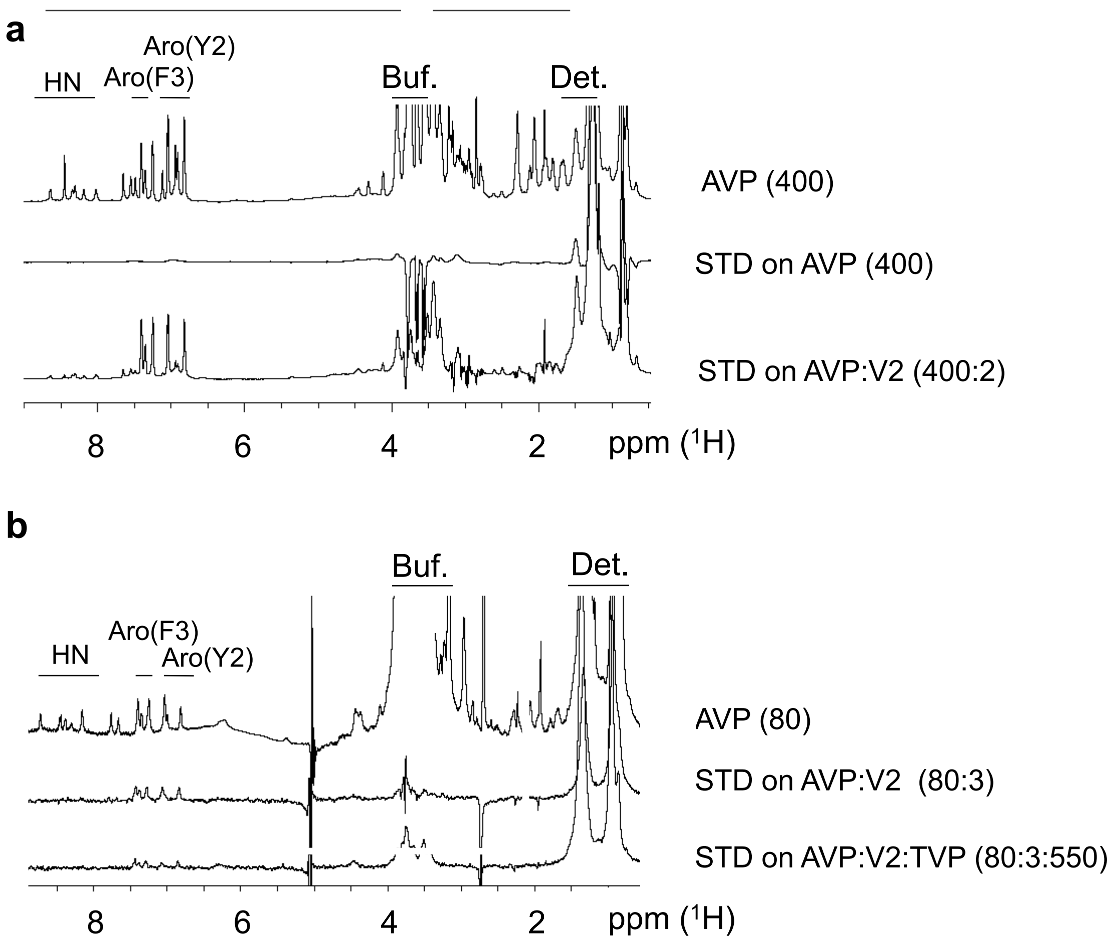

**Supplementary Figure 9: Principal components analysis (PCA) obtained from molecular dynamics simulation of the L state**

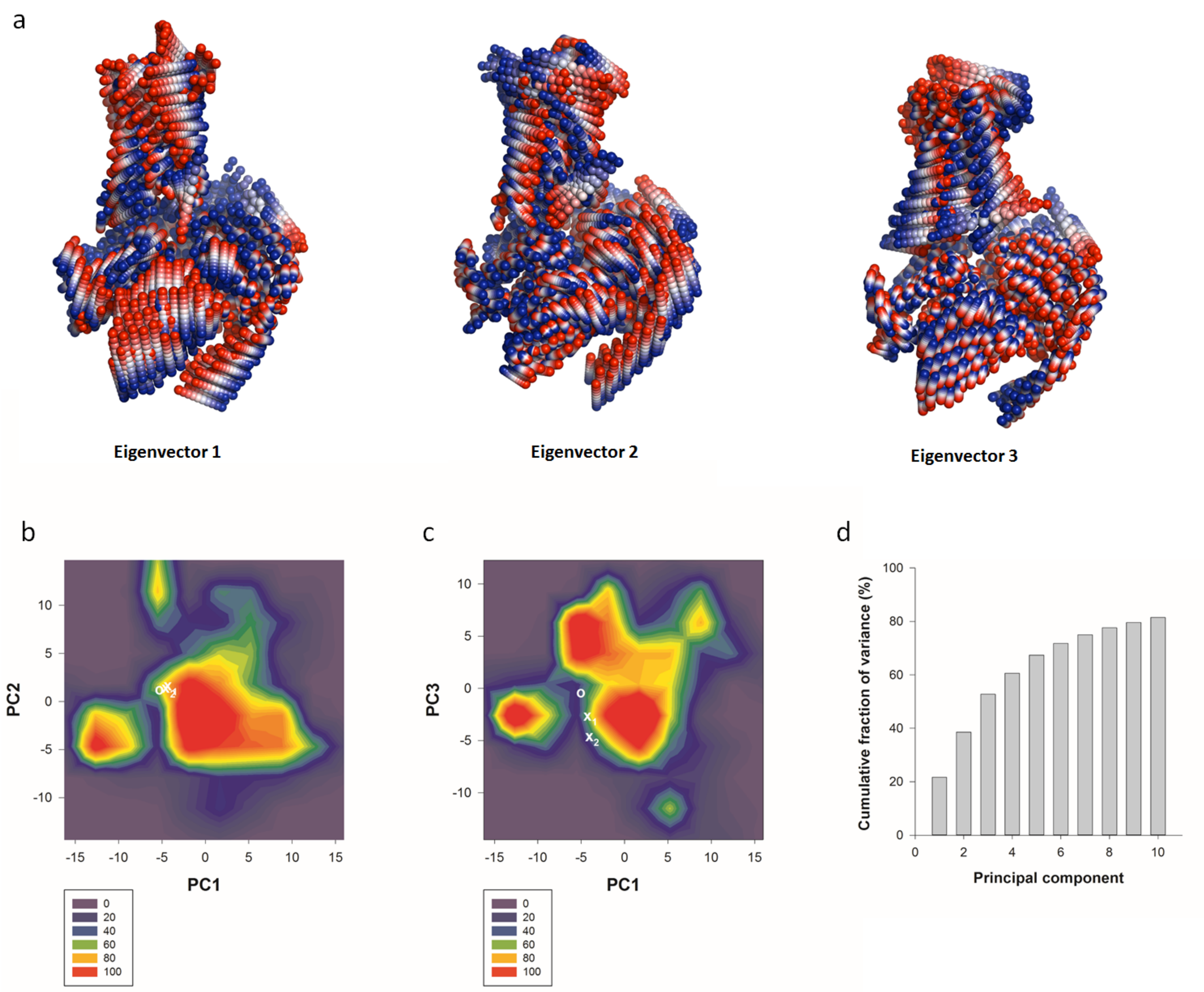

**Supplementary Figure 10 | Activation motifs and coupling interfaces in the AVP-V2R-Gs-Nb35 complex with the details of the cryo-EM maps**

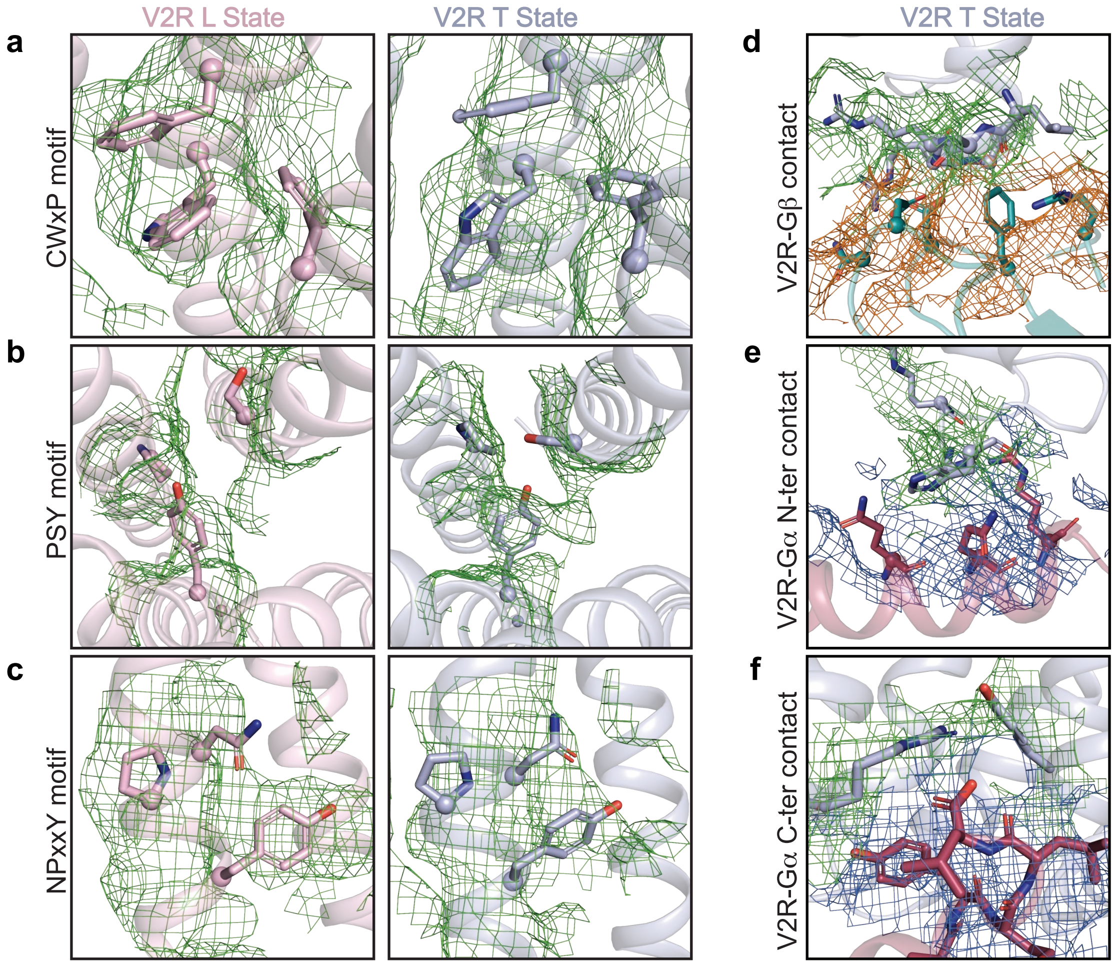
